# Pan-genome analyses of peach and its wild relatives provide insights into the genetics of disease resistance and species adaptation

**DOI:** 10.1101/2020.07.13.200204

**Authors:** Ke Cao, Zhen Peng, Xing Zhao, Yong Li, Kuozhan Liu, Pere Arus, Gengrui Zhu, Shuhan Deng, Weichao Fang, Changwen Chen, Xinwei Wang, Jinlong Wu, Zhangjun Fei, Lirong Wang

**Affiliations:** The Key Laboratory of Biology and Genetic Improvement of Horticultural Crops (Fruit Tree Breeding Technology), Ministry of Agriculture, Zhengzhou Fruit Research Institute, Chinese Academy of Agricultural Sciences, Zhengzhou 450009, China; Novogene Bioinformatics Institute, Beijing, P.R. China; IRTA, Centre de Recerca en Agrigenòmica, CSIC-IRTA-UAB-UB, Campus UAB – Edifici CRAG, Cerdanyola del Vallès (Bellaterra), Barcelona, Spain; Boyce Thompson Institute for Plant Research, Cornell University, Ithaca, NY 14853, USA; USDA-ARS, Robert W. Holley Center for Agriculture and Health, Ithaca, NY 14853, USA

**Keywords:** Peach, high-altitude adaptation, nitrogen recovery, pan-genome, non-linear evolution

## Abstract

As a foundation to understand the molecular mechanisms of peach evolution and high-altitude adaptation, we performed *de novo* genome assembling of four wild relatives of *P. persica, P. mira, P. kansuensis, P. davidiana* and *P. ferganensis*. Through comparative genomic analysis, abundant genetic variations were identified in four wild species when compared to *P. persica*. Among them, a deletion, located at the promoter of *Prupe.2G053600* in *P. kansuensis*, was validated to regulate the resistance to nematode. Next, a pan-genome was constructed which comprised 15,216 core gene families among four wild peaches and *P. perisca*. We identified the expanded and contracted gene families in different species and investigated their roles during peach evolution. Our results indicated that *P. mira* was the primitive ancestor of cultivated peach, and peach evolution was non-linear and a cross event might have occurred between *P. mira* and *P. dulcis* during the process. Combined with the selective sweeps identified using accessions of *P. mira* originating from different altitude regions, we proposed that nitrogen recovery was essential for high-altitude adaptation of *P. mira* through increasing its resistance to low temperature. The pan-genome constructed in our study provides a valuable resource for developing elite cultivars, studying the peach evolution, and characterizing the high-altitude adaptation in perennial crops.

---

Peach (*Prunus persica*) is the third most produced fruit crop, and is widely cultivated in temperate and subtropical regions. Due to its small genome size, peach has been used as a model plant for comparative and functional genomic researches of the Rosaceae family (Abbott et al., 2002). In 2013, a high-quality reference genome sequence of peach constructed with the Sanger whole-genome shotgun approach was released (International Peach Genome Initiative, 2013). Based on this genome sequence, researchers have investigated peach evolution (Cao et al., 2014b; Yu et al., 2018) and identified the domestication regions (Cao et al., 2014b; Akagi et al., 2016; Li et al., 2019) and genes associated with important traits (Cao et al., 2016; Cao et al., 2019). It is well known that wild germplasm contributes a significant proportion of the genetic resources of major crop species (Zhang et al., 2017), and significant phenotypic differences in fruit size, flavor, and stress tolerance were found among *P. persica* and its wild relatives, *P. mira, P. davidiana, P. kansuensis*, and *P. ferganensis* (Wang et al., 2012a). It is necessary to study genetic variations of peach and its wild relatives from a broader perspective, such as pan-genome analyses which have been conducted in other crops such as soybean (Li et al., 2014; Liu et al., 2020), rice (Wang et al., 2018; Zhao et al., 2018), sunflower (Hübner et al, 2019), tomato (Gao et al., 2019), etc. For example, after construction of a pan-genome of *Glycine soja*, Li et al. (2014) inferred that the copy number variations of resistance (R) genes could help to explain the resistance differences between wild and cultivated accessions.

Moreover, peach is an attractive model for studying high-altitude adaptability of perennial plants because its ancestral species, *P. mira*, originated in the Qinghai-Tibet plateau in China. The region has an average elevation of ∼4,000 m above the sea level, and the oxygen concentration is ∼40% lower and UV radiation is ∼30% stronger than those at the sea level (Yang et al., 2017). Up to date, knowledge on the mechanism of high-altitude adaptability has been reported in pig (Li et al., 2013), yak (Qu et al., 2013), human (Huerta-Sánchez et al., 2014; Yang et al., 2017), snakes (Li et al., 2018), hulless barley (Zeng et al., 2015), and the herbaceous plant *Crucihimalaya himalaica* (Zhang et al.,2019). However, little is known in perennial crops about the genetic basis of response to harsh conditions, such as low temperature and high UV radiation in high-altitude environments.

In the present study, we aimed to gain an in-depth understanding of the peach evolution and dissect genomic characteristics of some important agricultural traits. We *de novo* assembled the genomes of four wild relatives of *P. persica* to detect genomic variations and constructed a pan-genome of peach. Comparative genomic analysis identified a non-linear evolution event during peach evolution, and comprehensive characterization of selective sweeps revealed the mechanisms underlying the high-altitude adaptability in *P. mira*. Our study provides new insights into the peach evolution and help to dissect the genetic mechanism of important traits and to understand the interaction between perennial plants and climate from a genomic perspective.

## Results

### Assembly and annotation of unmapped reads of *P. persica*

Complete identification of genes in the *P. persica* genome is helpful to construct a sufficiently accurate pan-genome of peach. Therefore, we first sequenced 100 accessions belonging to *P. persica* with an average depth of 48.8× (Supplementary Table 1, Accession 1-100). An average of 3.4% of reads in each accession (Supplementary Table 1) failed to be aligned to the reference genome (Verde et al., 2017), and these unaligned reads were *de novo* assembled (Supplementary Table 2), which generated a total of 2.52-Mb sequences consisting of 2,833 non-redundant contigs (>500 bp) and a total of 923 non-reference (novel) genes (Supplementary Tables 3 and 4).

**Table 1.** Genomic sequencing, assembly, and annotation statistic for four wild peach species.

| Species | *P. mira* | *P. davidiana* | *P. kansuensis* | *P. ferganensis* |
| --- | --- | --- | --- | --- |
| Estimated genome size (Mb) | 242.94 | 237.29 | 238.06 | 237.24 |
| Sequencing depth (×) | 1350.58 | 263.69 | 120.76 | 126.88 |
| Total sequencing data (Gb) | 328.11 | 62.57 | 32.02 | 30.09 |
| Assembled genome size (Mb) | 252.39 | 220.51 | 206.19 | 204.58 |
| Contig N50 (bp) | 443,700 | 58,861 | 63,808 | 55,436 |
| Contig N50 (Number) | 159 | 940 | 838 | 1,031 |
| Scaffold N50 (bp) | 27,439,069 | 643,492 | 337,811 | 229,209 |
| Scaffold N50 (Number) | 4 | 87 | 150 | 228 |
| GC content (%) | 37.52 | 37.41 | 37.40 | 37.39 |
| Percent of repeat sequence (%) | 47.43 | 40.46 | 38.41 | 38.44 |
| Predicted gene number | 28,943 | 26,527 | 26,297 | 27,431 |
| Annotated gene number | 27,174 | 25,027 | 24,776 | 25,561 |

Combined with the reference genes (26,873), the total number of genes in the *P. persica* pan-genome was 27,796 (Supplementary Tables 3), among which 27,774 (99.92%) could be detected in the 100 resequenced accessions. According to the presence frequencies of detected genes in these accessions (Fig. 1a), we categorized them into core genes (24,971, 89.9%) that were shared by all the 100 accessions, and dispensable genes (2,803, 10.1%) that were defined as present in less than 99% of the accessions (Fig. 1b). The latter also can be divided into 356 softcore, 2366 shell and 81 cloud genes according to their presence frequencies higher than 99%, 1-99% and less than 1% of 100 peach accessions, respectively (Fig. 1b). Analyzing the relationship between the pan-genome size and iteratively random sampling accessions suggested a closed pan-genome with a finite number of both dispensable and core genes (Fig. 1c). In addition, the total and dispensable gene counts were obtained in different populations (Supplementary Fig. 1, Fig. 1d). As expected, of the 699 dispensable genes which were classified to be deficient in ornamental and wild *P. persica*, 59 were related to response to abiotic and biotic stress including those encoding NBS-LRR (nucleotide binding site-leucine-rich repeat) proteins. The enrichment of resistance (*R*) genes among the dispensable gene set was also observed in rice (Zhao et al., 2018). We also found that four genes in the dispensable gene set encoding geraniol 8-hydroxylases involved in terpene biosynthesis showed several tandem repeats at two loci in the Chr1: 24.47-24.66 Mb region. For example, *Prupe.1G231000* was detected in 63% of ornamental and wild *P. persica*, 88% of landraces, and 91% of improved varieties, indicating that this locus could be under positive selection during both domestication and improvement. This result may explain the rich terpene substances, such as linalool content, in improved vareities than that of landraces (Supplementary Fig. 2).

**Figure 1.**
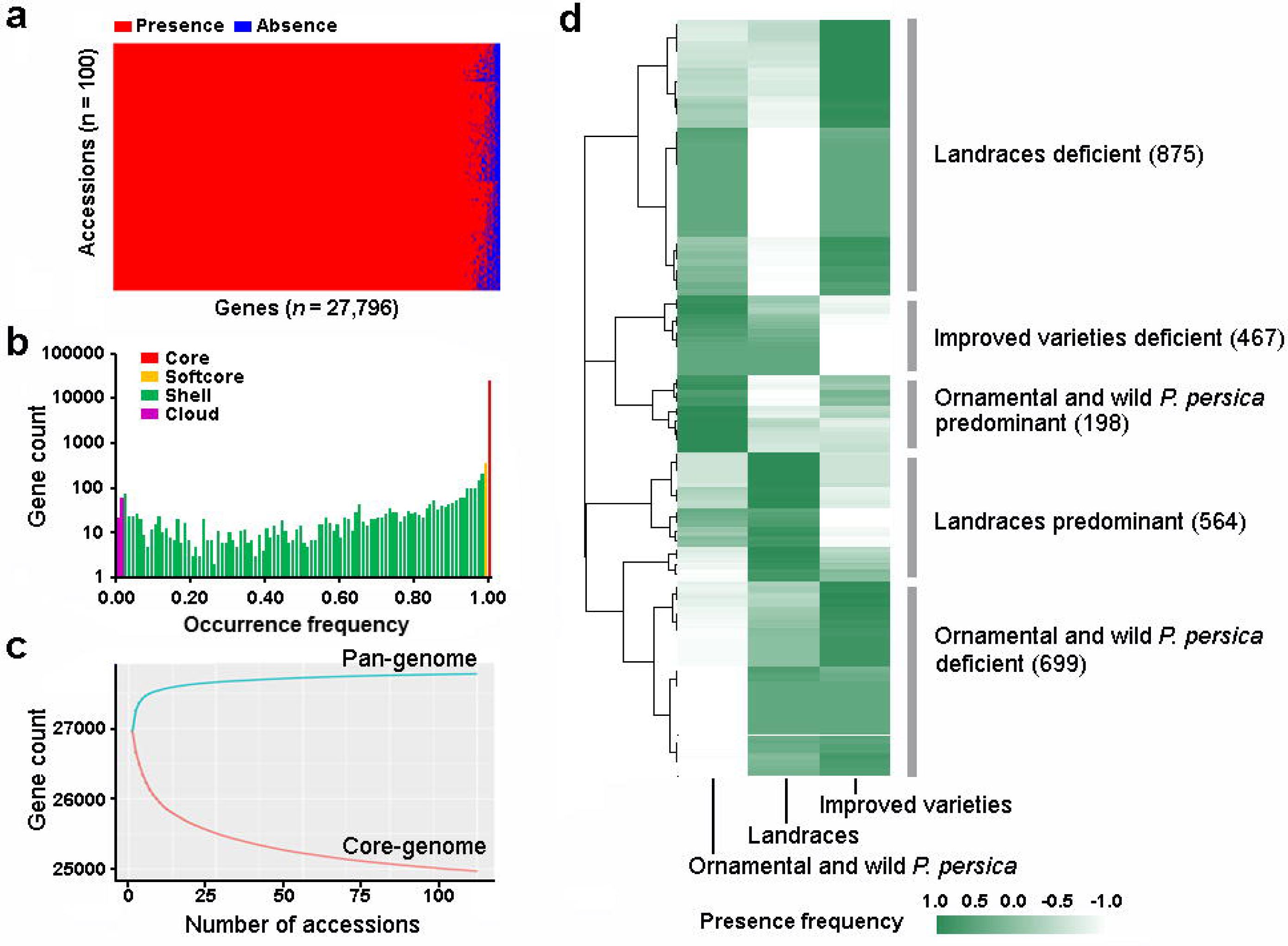
Pan-genome of *P. persica*. (a) Presence and absence information of genes in the 100 *P. persica* genomes. (b) Presence frequency of genes from the pan-genome of *P. persica*. (c) Simulation of the pan-genome and core-genome sizes in *P. persica*. (d) Presence frequency of 2,803 dispensable genes in the pan-genome of *P. persica* in different populations.

**Figure 2.**
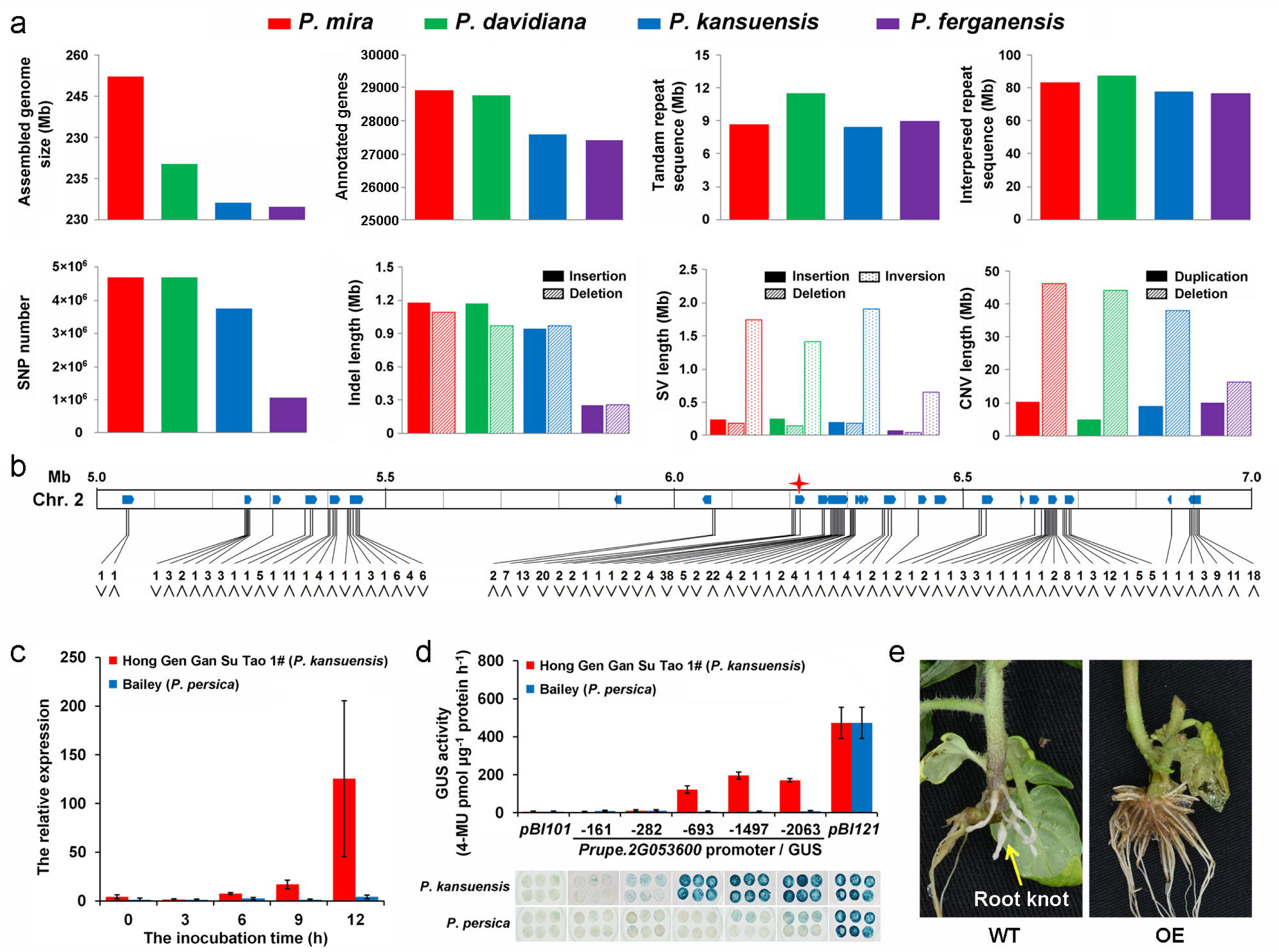
Variations identified in genomes of four wild peaches. (a) Genome size, annotated gene number, and the length of tandam repeat sequences, interpersed repeat sequences, and SNPs, small indels, SVs, and CNVs in *P. mira, P. davidiana, P. kansuensis*, and *P. ferganensis* compared to *P. persica*. (b) Variations in *P. kansuensis* but not in other wild peach species in the promoter and mRNA regions of annotated NBS-LRR genes on Chr. 2. “∧” indicates an insertion, “∨” indicates a deletion, and the asterisk indicates the candidate gene. (c) Expression of *Prupe.2G053600* in two accessions (‘Hong Gen Gan Su Tao 1#’ and ‘Bailey’) inoculated with nematode. (d) Promoter activity assay. Promoters with different lengths were fused to the *GUS* gene in plasmid pBI101. GUS was dyed and and its activity was measured using protein extracts of tobacco. N: negative control (pBI101), P: positive control (pBI121). (e) Functional validation of *Prupe.2G053600* through analysis of transgenic tomato plants expressing *Prupe.2G053600* under nematode treatment.

Moreover, we performed RNA-Seq analysis with different tissues (Supplementary Fig. 3, Supplementary Tables 5) as well as Sanger sequencing (Supplementary Fig. 4, Supplementary Tables 6), and confirmed that the novel sequences we assembled were reliable.

**Figure 3.**
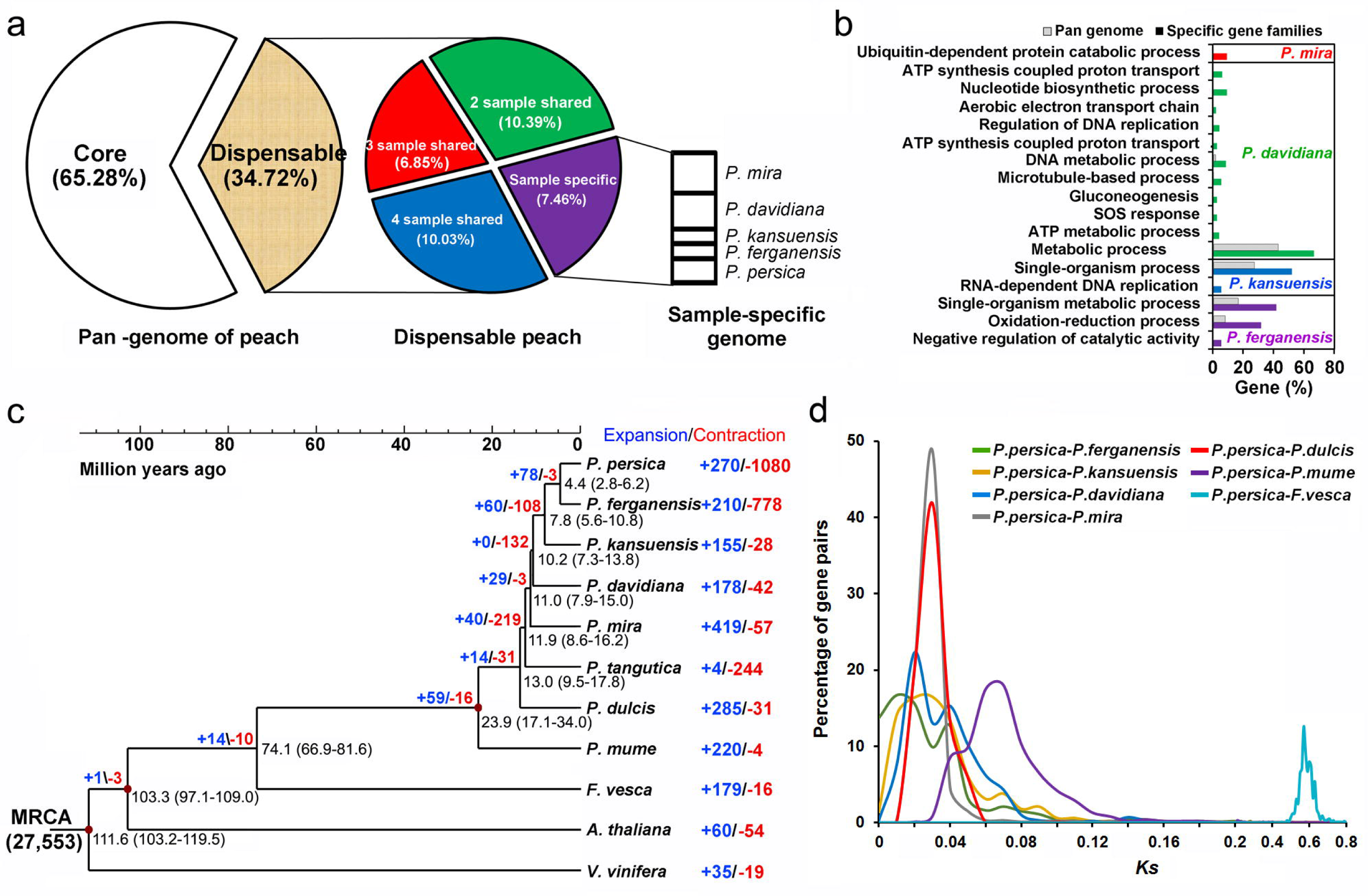
Pan-genome construction and evolutionary analysis of peach genome. (a) Core and dispensable gene families of four wild peaches and *P. persica*. (b) Gene Ontology annotation of genes specific in each species. (c) Estimation of divergence times of 11 species and identification of gene family expansions and contractions. Numbers on the nodes represent the divergence times from present (million years ago, Mya). MCRA, most recent common ancestor. (d) Distribution of *Ks* (synonymous mutation rate) values of orthologous genes between six genomes of the *Prunus* species and strawberry.

**Figure 4.**
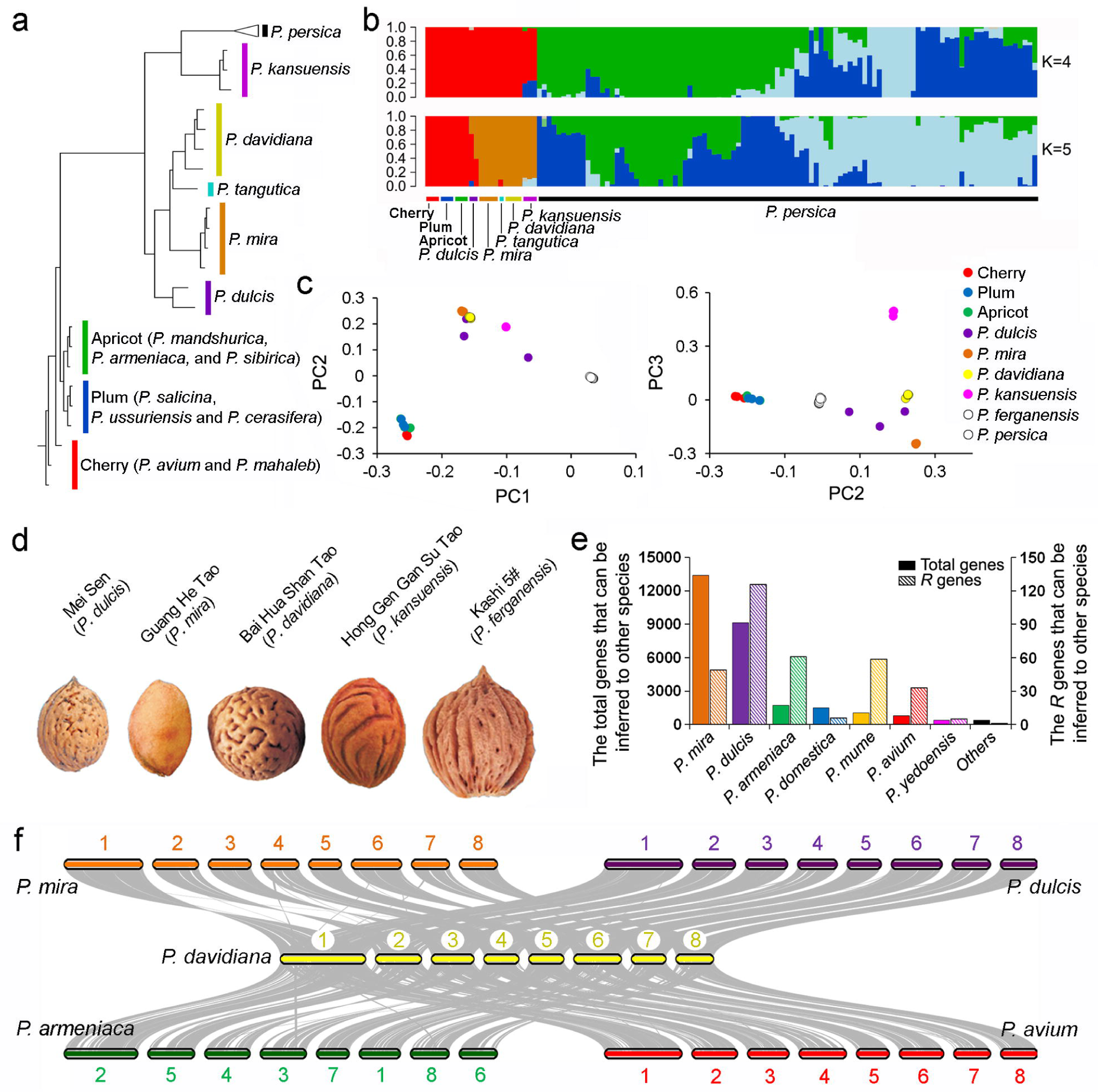
Nonlinear evolution of *P. davidiana*. (a) Phylogenetic tree, (b) the population structure, and (c) principal component analysis (PCA) of 126 peach accessions. (d) Stone steak morphology of *P. dulcis, P. mira, P. davidiana, P. kansuensis*, and *P. ferganensis*. (e) Statistics of gene pairs between *P. davidiana* and other *Prunus* species. (f) Collinearity between *P. davidiana* and *P. mira* or *P. dulcis, P. armeniaca*, and *P. avium* genomes.

### Assembly of the genomes of four wild peach species

The high-quality genome of *P. mira*, reckoned as the primitive of *P. persica* (Cao et al., 2014), was assembled using a more than 100-years old tree (Accession 123, Supplementary Fig. 5) through a combination of PacBio, Illumina, and Hi-C (High-throughput chromosome conformation capture) platforms. After estimating the genome size using the k-mer method (Supplementary Table 7, Supplementary Fig. 6), a total of 597.0× coverage of sequences were generated and used for genome assembly (Supplementary Table 8). A total of 657 scaffolds were anchored and 93.4% of them were allocated to eight pseudochromosomes (Supplementary Table 9). The contig and scaffold N50 sizes of the final assembly were 443.7 kb and 27.44 Mb, respectively (Table 1), which were higher than that of *P. persica* (255.42 kb and 27.37 Mb) sequenced using the Sanger technology (Verde et al., 2017).

**Figure 5.**
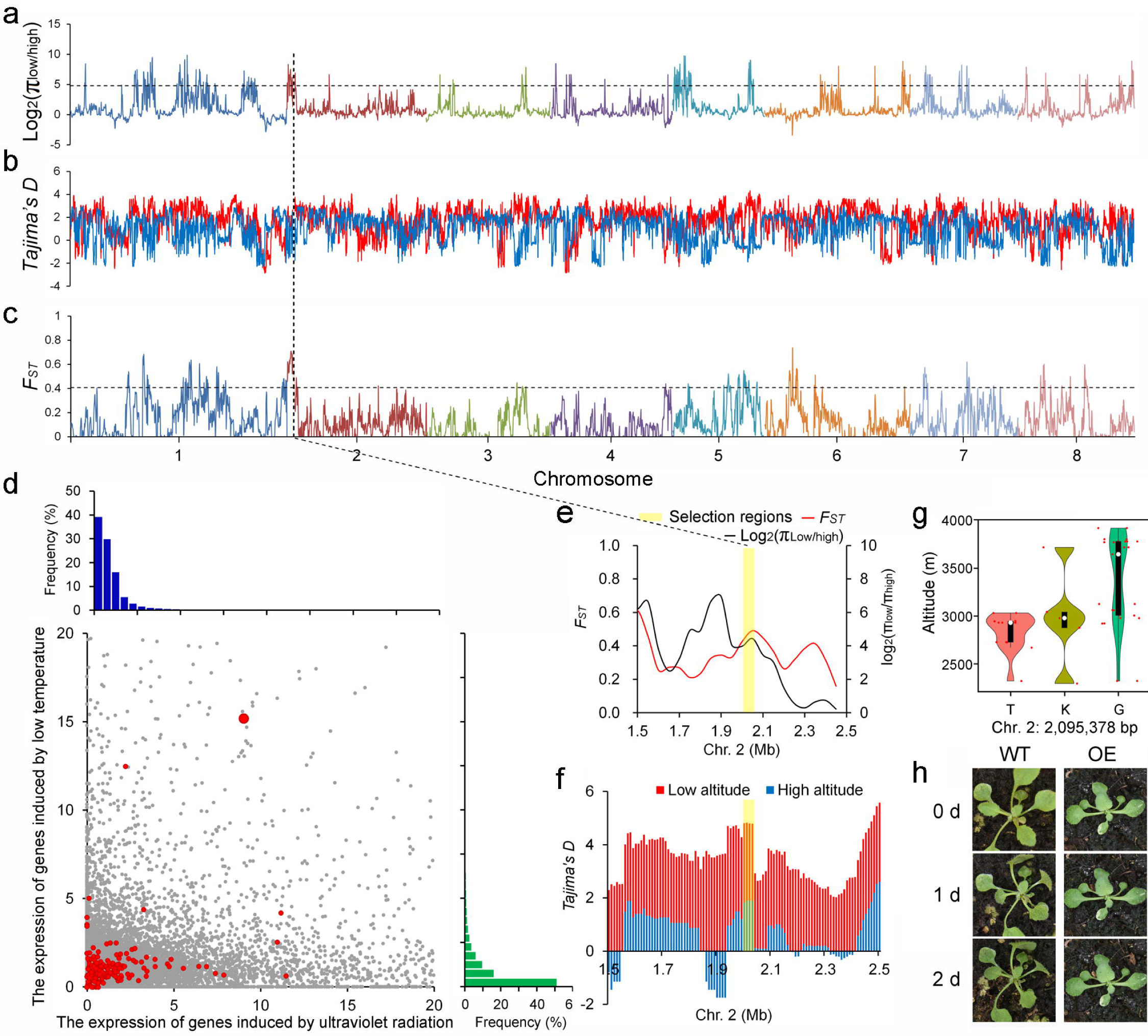
Selective regions associated with high-altitude adaptation in *P. mira*. (a-c)Domestication signals in accessions originating in high-altitude region compared to those in low-altitude. The signals were defined by the top 5% of π_ratio_ (a), Tajima’s D (b) and *F*_ST_ values (c). (d) Distribution of expression of genes induced by low temperature and UV of *P. mira*. Grey dots indicate the background genes and red dots indicate selective genes associated with high-altitude adaptation. (e) Detailed π ratio and *F*_*ST*_ values in the genome region of the candidate gene, *evm.model.Pm02.401* (pointed by the dashed line), which was substantially induced by low temperature. (f) Detailed Tajima’s D in the genome region of the candidate gene *evm.model.Pm02.401*. (g) Genotypes (K indicates G/T) of a variation (Chr. 2: 2,095,378 bp) located at the promoter of *evm.model.Pm02.401* in accessions from different altitude regions. (h) *A. thaliana* plants expressing *evm.model.Pm02.401* gene (OE) and the control (WT) treated with low temperature.

Draft genomes of three other wild peach species, *P. davidiana* (Accession 126), *P. kansuensis* (Accession 124), and *P. ferganensis* (Accession 125), were generated using only Illumina sequencing reads (Supplementary Table 8). We ultimately obtained 220.5, 206.2, and 204.6 Mb assemblies (Fig. 2a), covering about 92.9%, 86.6%, and 86.2% of the estimated genome sizes and having the scaffold N50 lengths of 0.64, 0.34, and 0.23 Mb for *P. davidiana, P. kansuensis*, and *P. ferganensis*, respectively (Table 1). The high quality of the assemblies was demonstrated using the BUSCO (Simao et al., 2015) analysis (Supplementary Table 10) and RNA-Seq read mapping rates (Supplementary Table 11).

An overview of the genome synteny between *P. mira* and *P. persica* is presented in Supplementary Fig. 7. As found in other plant genomes, long terminal repeat (LTR) retrotransposons made up the majority of the transposable elements (TEs), comprising about 25.4% of the *P. mira* genome. In contrast with *P. persica, P. mira* had a higher percentage of DNA transposons in the genome (Supplementary Table 12; 9.1% in *P. persica* vs 15.0% in *P. mira*). Subsequently, gene prediction and annotation were performed resulting in 28,943, 26,527, 26,297, and 27,431 protein-coding genes in *P. mira, P. davidiana, P. kansuensis*, and *P. ferganensis*, respectively (Fig. 2a, Supplementary Table 13 and 14). We found substantially lower densities of repeat sequence as well as higher gene density near the telomeres of each chromosome (Supplementary Fig. 7a, b). The accumulated gene expression level was higher in regions with higher gene densities (Supplementary Fig. 7c-g). About 93.2-94.3% of the protein-coding genes of the four wild species could be functionally annotated (Supplementary Table 15). In addition, we identified 49-195 ribosomal RNA, 476-541 transfer RNA, 340-449 small nuclear RNA, and 409-489 microRNA genes in the four wild peach species (Supplementary Table 16).

### A root-knot nematode resistance gene identified through genome comparison

To discover sequence variations, we anchored the four assembled wild genomes onto the reference genome of *P. persica* (Verde et al., 2017). A total of 1,062,698-4,683,941 single nucleotide polymorphisms (SNPs; Supplementary Table 17), 157,379-691,686 small insertions and deletions (indels; Supplementary Table 18), 2,475-8,418 large structural variants (SVs including insertions, inversions, and deletions; ≥ 50 bp in length; Supplementary Table 19), and 4,153-7,090 copy number variations (CNVs including deletions and duplications; Supplementary Table 20) were identified in the four species (Fig. 2a, Supplementary Fig. 8). It was unexpected that *P. davidiana* had more SNPs, insertions of SVs, and deletions of CNVs than *P. mira* because the latter was recognized as the oldest ancestor of *P. persica* harboring a longer genetic distance with *P. persica* (Cao et al., 2014b).

Based on the variation detection, we found that an obvious positive selection existed during evolution according to the ratio of the number of nonsynonymous to synonymous SNPs (1.26, 1.24, 1.28, and 1.50 in *P. mira, P. davidiana, P. kansuensis, P. ferganensis*, respectively) and non-frameshift indels to frameshift ones (1.43, 1.46, 1.46, and 2.22 in *P. mira, P. davidiana, P. kansuensis, P. ferganensis*, respectively) in CDS in different species. Next, we analyzed the function of the genes which comprised different variations (Supplementary Fig. 9-12) and found that plant-pathogen interaction pathways were enriched in genes containing small indels and CNVs in all four wild relatives of peach.

Analysis of genomic variations through pan-genome analysis allowed us to identify candidate genes involved in important agronomic traits. Root-knot nematodes are an important pest that seriously damages peach. We previously constructed a BC_1_ population from the cross between ‘Hong Gen Gan Su Tao 1#’ (*P. kansuensis*) and a cultivated peach ‘Bailey’ (*P. persica*). ‘Hong Gen Gan Su Tao 1#’ harbored high resistance to root-knot nematodes (*Meloidogyne incognita*), whereas other accessions used for pan-genome construction including *P. mira, P. davidiana*, and *P. ferganensis*, all showed low resistance to *M. incognita* (Zhu et al., 2000). Using this BC_1_ population, a nematode resistance locus was mapped at the top region of Chr. 2 (5.0-7.0 Mb) (Cao et al., 2014a). In other species, the R genes to nematodes generally encode proteins containing the NBS-LRR domain (Cao et al., 2014a). The genome variations mainly small indels and SVs were then compared in different species and 78 of them, which only occurred in the gene and promoter regions in *P. kansuensis* but not in other species, were identified in the nematode resistance locus on Chr. 2. Among them, 24 were annotated as R genes (Supplementary Table 22) and one of them, *Prupe.2G053600*, which comprised a large deletion in the promoter was further analyzed (Fig. 2b). qRT-PCR analysis revealed that the gene was differentially expressed in roots of ‘Hong Gen Gan Su Tao 1#’ and ‘Bailey’ innoculated with *M. incognita* (Fig. 2c). We validated this promoter deletion and found that it co-segregated with resistant phenotype of the seedlings in the BC_1_ population. To identify the active region of the promoter in *Prupe.2G053600*, we amplified 161, 282, 693, 1497, and 2063 bp of the 5′ flanking region of the gene in ‘Hong Gen Gan Su Tao 1#’ and linked the amplified products with the β-glucuronidase (GUS) coding sequence to transiently transformed into *Nicotiana tabacum*. Leaves from the transgenic lines were analyzed for GUS activity by histochemical GUS staining and GUS quantitative enzyme activity determination. The lines carrying the various *Prupe.2G053600* promoters displayed remarkable but lesser GUS activity in comparison with the CaMV35S transformed one (pBI121 vector). An increase in GUS expression was observed with promoters longer than 693 bp, indicating the deletion (310 bp ahead of start codon) could drive the expression of *Prupe.2G053600* (Fig. 2d). Finally, the coding sequence of this gene were inserted into the plant expression vectors and transformed into tomato (cv. Micro-Tom). The transgenic lines were validated by genomic PCR and qRT-PCR. One transgenic line with high expression of *Prupe.2G053600* was selected to analyze nematode resistance after 7 d post infection. We found the transgenic line showed remarkable nematode resistance with less root knots compared to control plants (Fig. 2e).

### Peach genome evolution and species divergence

Regarding the core and dispensable portions of the *P. mira, P. davidiana, P. kansuensis, P. ferganensis*, and *P. persica* genomes, all of the genes in the five genomes could be classified into 23,309 families on the basis of the homology of their encoded proteins (Supplementary Fig. 13a). The comparison of above species revealed 8,093 (34.7%) dispensable gene-families distributed across all genomes, and 543, 485, 194, 197, and 320 families specific to each of the above species, respectively (Fig. 3a). Ubiquitin-dependent protein catabolic process, metabolic process, single-organism process, and oxidation-reduction process were found to be enriched in gene families specific to *P. mira, P. davidiana, P. kansuensis*, and *P. ferganensis*, respectively (Fig. 3b).

In the gene families, we identified 3,548 single-copy orthologs in the four wild peaches (Supplementary Fig. 14). Using these single-copy orthologs, we constructed a phylogenetic tree of *P. persica* and its wild related species as well as other representative plant species (Fig. 3c). Based on the known divergence time between *Arabidopsis thaliana* and strawberry, the age estimate for the split of *P. mume* and the common ancestor of *P. persica* and its wild relatives was around 23.9 million years ago (Mya), later than that in a previous report, presumably 44.0 Mya (Baek et al., 2018). The divergence time of *P. dulcis* and *P. mira* was about 13.0 Mya, which was obviously earlier than that of Yu et al. (2018) and Alioto et al. (2020) who found the divergence time of the two species was 4.99 and 5.88 Mya, respectively. Furthermore, we found that *P. mira* split with the common ancestor of *P. davidiana, P. kansuensis*, and *P. ferganensis* approximately 11.0 Mya. The event occurred around the drastic crustal movement of Qinghai-Tibet Plateau (Chung et al., 1998) where *P. mira* originated.

To validate the speciation events of peach species, fourfold synonymous third-codon transversion (4DTv) rates were calculated for a total of 2,273, 2,746, 3,438, and 4,386 pairs of paralogous genes in *P. ferganensis, P. kansuensis, P. davidiana* and *P. mira*, respectively. We found that all 4DTv values (Supplementary Fig. 15) among paralogs in four wild peach species peaked at around 0.50 to 0.60, consistent with the whole-genome triplication event (γ event) shared by all eudicots and indicating that no recent whole-genome duplication occurred. A peak 4DTv value at around 0 for the orthologs between *P. persica* and *P. mira* highlighted a very recent diversification of *Prunus* species (Baek et al., 2018). To estimate the time of species divergence of the four wild species, we calculated the *Ks* (rate of synonymous mutation) values of orthologous genes between these species. As shown in Fig. 3d, the peaks at a *Ks* mode of 0.03 for orthologs between *P. persica*-*P. mira* and *P. persica*-*P. dulcis* genomes indicated similar divergence time of the *P. mira* and *P. dulcis* from *P. persica*, consistent with the results of the phylogenetic tree.

### A non-linear event involved in the genome evolution of *P. davidiana*

In the previous study, *P. mira* was recognized as the primitive of *P. persica* (Cao et al., 2014). In this study, we found that the SNPs, insertions of SVs, deletions of CNVs as well as tandem repeat sequences and *R* genes were more abundant in *P. davidiana* than in the other wild related species when compared with *P. persica* (Fig. 2a). Meanwhile, k-mer frequency distribution (two peaks) clearly indicated the high heterozygosity level of the *P. davidiana* genome (Supplementary Fig. 6b). In addition, two peaks were also found in *Ks* values of orthologous genes between *P. davidiana* and *P. persica*, and between *P. ferganensis* and *P. persica* (Fig. 3d). Therefore, we speculated that the evolution of *P. davidiana* was not linear and the novel sequences might come from a crossing event.

Twenty-six accessions of wild peach species (Supplementary Table 1, accessions 101-126) were resequenced, and the obtained sequences together with those from the 100 *P. persica* accessions were aligned to the *P. mira* genome to obtain a total of 839,431 high-quality SNPs. We performed phylogenetic (Fig. 4a) and structure analyses (Fig. 4b) and found that the most primitive species of peach was *P. mira*, followed by *P. davidiana* and *P. kansuensis*, similar to those reported in a previous study (Yu et al., 2018) that *P. tangutica* and *P. davidiana* were closely related and *P. dulcis* first differentiated from the *Persica* section of subg. *Amygdalus*. However, principal component analysis (PCA) demonstrated that *P. dulcis* located between *P. mira* and *P. davidiana* in the diagram of PC2-PC3 (Fig. 4c). We then phenotyped the stone streak of different species and found that a lot of dot streaks were present in the *P. davidiana* but absent in the more recent species, *P. kansuensis*. However, the dot streaks were found in the ancient species of this family, such as *P. dulcis* (Fig. 4d). To further validate the nonlinear event that occurred during peach evolution from *P. mira* to *P. persica* and identify the potential parent of *P. davidiana*, all the genes in *P. davidiana* were aligned with those from the putative ancestors (Fig. 4e), including *P. mira* and *P. dulcis*. The genes were then assigned as originating from a specific species if the highest score was observed from the alignments of orthologous genes between *P. davidiana* and the species. The results showed that about 47.5% of the genes (13,369) were specific to *P. mira*, 32.3% (9,081) to *P. dulcis*, and no more than 6% to other species.

According to the above evidence, we propose that *P. dulcis* is ancestral to *P. mira, P. davidiana* and others, same as that of Yu et al. (2018). However, *P. davidiana* showed intermediate genomic characteristics between *P. mira* and *P. dulcis* and might originate from the cross between these two species. This finding is different from the previous study which focused on an ancient introgression between *P. mira* and the common ancestor of *P. kansuensis* and *P. persica* (Yu et al., 2018). Meanwhile, since no reference genome is available for *P. tangutica*, direct comparison of genes from *P. davidiana* and *P. tangutica* is not feasible. Therefore, we first aligned the sequences of *P. davidiana* to its putative ancestor, *P. mira*, and assembled the unmapped sequences to obtain a partial reference genome. Genome resequencing data of different species were then aligned to this partial genome. We found that *P. dulcis*, not *P. tangutica*, showed the highest mapping rates (Supplementary Fig. 16), which again proved that the introgression in *P. davidiana* came from *P. dulcis* although *P. tangutica* has a closer relationship with *P. davidiana* (Fig. 4a). To further study the evolutionary events leading to the genome structure of *P. davidiana*, we connected all assembled contigs of *P. davidiana* to pseudochromosomes using the Hi-C technology and investigated the chromosome-to-chromosome relationships based on 153 (*P. davidiana* versus *P. mira*) and 125 (*P. davidiana* versus *P. dulcis*) identified syntenic blocks (Fig. 4f). The mosaic syntenic patterns again demonstrated that *P. davidiana* might have arisen during the evolutionary process of *P. mira* but with a cross of *P. dulcis*. Although *P. dulcis* is mainly distributed in Georgia, Azerbaijan, Turkey, Syria and Xinjiang province (China), it can also be found in Sichuan province of China where some *P. mira* grow in this region at the same time, indicating the hybridization event is highly possible (Supplementary Fig. 17).

### Genomic basis of pathogen resistance in peach

Plants have to face various biotic and abiotic stresses during their growth and development. Among *P. persica* and its four wild relatives, *P. davidiana* is widely distributed in northern China, while others such as *P. mira, P. kansuensis*, and *P. ferganensis* grow only in one specific region, such as Qinghai-Tibet Plateau, Gansu, and Xinjiang province of China, respectively. How these species respond to different geographical environments at the genomic level remains unclear.

*R* genes are of particular interest because they confer resistance against a series of pests and pathogens. In this study, a total of 310, 339, 323, and 320 putative *R* genes were identified in *P. mira, P. davidiana, P. kansuensis*, and *P. ferganensis* genomes, respectively (Supplementary Table 23). The largest number of *R* genes in *P. davidiana* might explain its strong and multiple resistances to different pathogens, such as aphid, *Agrobacterium tumefaciuns*, etc. In addition, the least R genes identified in *P. mira* might be due to few pathogenic infections in the Qinghai-Tibet Plateau with a cold weather and strong ultraviolet light environment, similar to contracted *R* gene family observed in *Crucihimalaya himalaica* also with a typical Qinghai-Tibet Plateau distribution (Zhang et al. 2019). We found that R genes were distributed across the eight chromosomes unevenly in all four wild peaches (Supplementary Fig. 18), similar to the findings in pear (Wu et al., 2013), kiwifruit (Huang et al., 2013) and jujube (Liu et al., 2014). We compared the previously identified disease-resistance QTLs/genes with the distribution of R genes and found most of the QTLs/genes were located in genome regions containing candidate *R* genes (Supplementary Fig. 19).

We further analyzed the origin of R genes in *P. davidiana*, which harbored the largest number of R genes among the four wild peaches. Of all 339 R genes in *P. davidiana*, 37.2% were categorized to originate from *P. dulcis*, followed by *P. armeniaca* (18.0%), and *P. mume* (17.4%), while only 49 (14.5%) from *P. mira* (Fig. 4e). Therefore, we hypothesize that the cross between *P. mira* and *P. dulcis* enhanced the adaptation of *P. davidiana* when it was spread to new environments.

### Genomic basis of adaptation to high altitude in *P. mira*

Analysis of the adaptation of *P. mira* to high altitude is helpful to discover genes or loci that can be used in breeding programs to expand the cultivation area of peach. When analyzing the genome variations in different species, we found that genes comprising small indels (Supplementary Fig. 10) and large SVs (Supplementary Fig. 11) were both enriched with those related to purine metabolism. Furthermore, lineage-specific gene family expansions may be associated with the emergence of specific functions and physiology (Kim et al., 2011). With the genome evolution of wild peach relatives, the number of expanded or contracted gene families were decreased from *P. mira* to *P. kansuensis* while increased in *P. ferganensis* and *P. persica* compared with that in the most recent common ancestor (MCRA, Fig. 3c; Supplementary Fig. 13b and 13c). We found that the expanded gene families were alo highly enriched with those related to purine metabolism in *P. mira* (Supplementary Fig. 19), same as in *C. himalaica* which grows in the same regions (Zhang et al., 2019). Further analysis indicated that among the above gene families, a total of 225 genes encoding (*S*)-ureidoglycolate amidohydrolase (UAH) were identified in *P. mira* and the corresponding gene numbers decreased to 5, 4, 2, and 2 in *P. davidiana, P. kansuensis, P. ferganensis*, and *P. persica*, respectively. Enzymes encoded by this gene family catalyze the final step of purine catabolism, converting (*S*)-ureidoglycolate into glyoxylate (Werner et al., 2010). It is well known that nitrogen recycling and redistribution are important for plants responding to the environmental stresses, such as drought, cold, and salinity (Alamillo et al., 2010; Kanani et al., 2010; Yobi et al., 2013). Interestingly, *OsUAH* has been identified as being regulated by low-temperature in rice, and a C-repeat/dehydration-responsive (CRT/DRE) element in its promoter specifically binds to a C-repeat-binding factor/DRE-binding protein 1 (CBF/DREB1) subfamily member, OsCBF3, indicating its function in low temperature tolerance (Li et al., 2015). Therefore, the enrichment of *UAH* genes in *P. mira* might explain its high-altitude adaptability.

Population genomic analyses were also performed to analyze the high-altitude adaptability of *P. mira*. A total of 32 accessions (Supplementary Table 1) belonging to the species with an altitude ranging from 2,290 to 3,930 m were resequenced to an average depth of 40.1×, and the sequencing reads were aligned to the *P. mira* genome to identify a total of 1,394,483 SNPs. Based on the phylogenetic and structure analyses using the identified SNPs, three accessions (Linzhi 8#, Guang He Tao 27#, and Guang He Tao 57#) thought to have not corresponded to their altitude categories, two (Guang He Tao 29# and Guang He Tao 50#) reckoned as an admixture subgroup between high and low altitude subgroups, and one (Guang He Tao 28#) showing large genetic distance with others were excluded in the downstream analysis. Then, six accessions were classified into a high-altitude subgroup and 20 into the low-altitude subgroup (Supplementary Fig. 20). We calculated and compared the nucleotide diversity (π; Fig. 5a), Tajima’s *D* (Fig. 5b), and *F*_*ST*_ (Fig. 5c) values using SNPs across the genome of high- and low-altitude groups, resulting in the identification of selective sweeps of a total of 789 kb and containing 222 genes (Supplementary Table 24). These genes were mostly involved in resistance to a series of stresses, such as cold, UV light, and DNA damage (Supplementary Table 25, Supplementary Fig. 21). Furthermore, using the young seedlings of *P. mira* treated under low temperature and UV-light for 10 h, we found that most genes in the selective sweeps presented stronger induction by UV than by low temperature based on the RNA-Seq data (Fig. 5d). However, one gene, *evm.model.Pm02.401*, encoding a CBF/DREB1 protein, showed more than 3,000-fold induction of expression by cold and about 60-fold induction by UV-light. According to the *F*_*ST*_ and Tajima’s D values, this gene was indeed under selection by altitude (Fig. 5e, 5f). Based on resequencing data, we found five SNPs showing a strong association with the phenotype, including a SNP located at 1,222 bp upstream of the start codon (Fig. 5g). In addition, we heterologously expressed the *evm.model.Pm02.401* gene in Arabidopsis, and the transgenic plants were exposed to 0 °C for 24 h. The transgenic Arabidopsis seedlings showed increased resistance to low temperature compared to the wild type (Fig. 5h). Together these results indicated the selection and expression of the *evm.model.Pm02.401* gene were associated with low temperature resistance of peach in high-altitude regions. Combined with the pervious study in rice (Li et al., 2015), we believe that CBFs/DREB 1s and its target genes in the UAH family may play important roles in plateau adaptability of *P. mira*.

### Discussion

In peach, a high-quality reference genome of *P. persica* was released and has since widely used as a valuable resource for effectively mining candidate genes for important traits (Verde et al., 2017). However, this genome sequence alone is not adequate to uncover wild-specific sequences which might have been lost during domestication or artificial selection (Xie et al., 2019). In this study, we first constructed a pan-genome of *P. persica*, with a total of 2.52-Mb non-reference sequences and comprising 923 novel genes. We then *de novo* assembled the genomes of four wild relatives of *P. persica*, including a species (*P. mira*) that exclusively originated in the Qinghai-Tibet Plateau. Using this large-scale comprehensive dataset, millions of genomic variations including SNPs and SVs were identified. Finally, a pan-genome of all peach species was constructed and hundreds of specific gene families in each of the wild peach species were identified. The above gene sets represent a useful source for in-depth functional genomic studies including the identification of a nematode resistance gene from *P. kansuensis* and the elucidation of the evolution history of *P. davidiana*. The nonlinear evolution of peach identified in this study expands our understanding of the evolutionary path of peach and plant speciation. In addition, based on expanded gene families and comparative genomic analysis using different accessions of *P. mira* originating from low- and high-altitude regions, a new mechanism underlying high-altitude adaptation in *P. mira*, high nitrogen recovery, was discovered. These findings provide important insights into the similarities and differences in high-altitude adaptive mechanisms among perennial, annual plants and animals.

## Methods

### Plant materials

In the study, different samples were used for DNA sequencing. First, 100 peach accessions belonging to *P. persica* were used for genome resequencing and construction of a pan-genome of *P. persica*. Second, genomes of four wild accessions (2010-138, Zhou Xing Shan Tao 1#, Hong Gen Gan Su Tao 1#, and Ka Shi 1#) were sequenced and *de novo* assembled. Third, 26 accessions from different wild peach species were selected for genome resequencing. The above accessions were conserved in the National Germplasm Resource Repository of Peach at Zhengzhou Fruit Research Institute, CAAS, China. Fourth, 32 accessions belonging to *P. mira* were sampled from Tibet with different altitudes and their genomes were resequenced. Fifth, one BC_1_ population was constructed between ‘Hong Gen Gan Su Tao 1#’ (*P. kansuensis*) and ‘Bailey’ (*P. persica*) to identify QTLs linked to nematode resistance. Resistance to nematode in this BC_1_ population was evaluated previously by our group (Cao et al., 2014a). Genomic DNA was extracted using the Plant Genomic DNA kit (Tiangen, Beijing, China) from young leaves.

Moreover, different samples were used for RNA sequencing (RNA-Seq). First, young leaves, mature fruits, seeds, phloem, and roots (obtained through asexual reproduction) of *P. persica* (Shang Hai Shui Mi), *P. ferganensis* (Kashi 1#), *P. kansuensis* (Hong Gen Gan Su Tao 1#), *P. davidiana* (Hong Hua Shan Tao) and *P. mira* (2010-138) were collected. Second, roots of ‘Hong Gen Gan Su Tao 1#’ and ‘Bailey’ infected with *Meloidogyne incognita* for 3, 6, 9, 12 h were collected for RNA-Seq analysis.

In addition, the mature fruits of 57 peach varieties were selected to evaluate linalool content using gas chromatograph-mass spectrometer after extracting volatile substances by headspace microextraction method in 2015 and 2016 (Luo et al., 2017).

### Pan-genome construction of *P. persica*

SOAPdenovo2 (Luo et al., 2012) was used to assemble the genomes of 100 *P. persica* accessions with *k*-mer set to 31. The quality of the genome assembly was assessed using QUAST (version 2.3) (Gurevich et al., 2013) with the peach reference genome (Verde et al., 2017). From QUAST output, unaligned contigs longer than 500 bp were retrieved and merged. CD-HIT (Fu et al., 2012) version 4.6.1 was used to remove redundant sequences with parameters ‘-c 0.9 -T 16 -M 50000’. For the remaining sequences, all-versus-all alignments with BLASTN were carried out to ensure that these sequences had no redundancy. Next, the non-redundant sequences were aligned to the GenBank nt database with BLASTN with parameters ‘-evalue 1e-5 -best_hit_overhang 0.25 -perc_identity 0.5 -max_target_seqs 10’. Contigs with the best alignments (considering *E*-values and identities) not from Viridiplantae or from chloroplast and mitochondrial genomes were considered as contaminants and removed. The remaining contigs formed the non-redundant novel sequences. The pan-genome of *P. persica* species was then generated by combining the reference peach genome and non-redundant novel sequences.

The non-redundant novel sequences were annotated with *ab initio*, homology-based and transcript-based predictions. Genome sequences of the 100 accessions were then mapped to the pan-genome, and based on the alignments the presence or absence of each gene in the pan-genome in each accession was inferred.

### Confirmation of the unmapped contigs

In order to verify the assembled contigs from the 100 *P. persica* accessions that were not mapped to the peach reference genome, we randomly selected 10 contigs for designing 10 pairs of primers. PCR were then performed to amplify these 10 contigs in 8 accessions belonging to different geographic groups. The resulting PCR products were sequenced using the Sanger technology and sequences were aligned to the template sequence with DNAman software.

### Genome sequencing of wild peach species

The *P. mira* (2010-138) genome was sequenced using different platforms including PacBio Sequel and Illumina, and the other species were sequenced only using Illumina platform, according to the manufacturers’ protocols. Library construction and sequencing was performed at Novogene Bioinformatics Technology Co., Ltd (Tianjin, China). For short-read sequencing, two short-insert libraries (230 bp and 500 bp) and 4 large-insert libraries (2 kb, 5 kb, 10 kb, and 20 kb) were constructed for *P. mira* and *P. davidiana*, while two short-insert libraries and 2 large-insert libraries were constructed for *P. kansuensis* and *P. ferganensis*. These libraries were sequenced on an Illumina HiSeq X Ten platform.

For single molecule real-time (SMRT) sequencing for *P. mira*, a 20-kb library was constructed and sequenced on the PacBio Sequel platform.

For Hi-C sequencing, leaves fixed in 1% (vol/vol) formaldehyde were used for library construction. Cell lysis, chromatin digestion, proximity-ligation treatments, DNA recovery and subsequent DNA manipulations were performed as previously described (Lieberman-Aiden, 2009). MboI was used as the restriction enzyme in chromatin digestion. The Hi-C library was sequenced on the Illumina HiSeq X Ten platform to generate 150 bp paired-end reads.

### RNA-Seq data generation

To assist protein-coding gene predictions, we performed RNA-Seq using five different tissues for each species, and for each sample, three independent biological replicates were generated. Total RNA was extracted with the RNA Extraction Kit (Aidlab, Beijing, China), following the manufacturer’s protocol. RNA-Seq libraries were prepared with the Illumina standard mRNA-seq library preparation kit and sequenced on a HiSeq 2500 system (Illumina, San Diego, CA) with paired-end mode.

### Genome assembly of wild peach species

The genome sizes of the four wild peach species were estimated by K-mer analysis. The occurrences of K-mer with a peak depth were counted using Illumina paired-end reads, and genome sizes were calculated according to the formula: total number of K-mers / depth at the K-mer peak, using JELLYFISH 2.1.3 software (Marcais and Kingsford, 2011) with K set to 17.

Illumina reads from the four wild species were assembled using ALLPATHS-LG (Butler et al. 2008), and gaps in the assemblies were filled using GapCloser V1.12 (Luo et al., 2012). Mate-paired reads were then used to generate scaffolds using SSPACE (Boetzer et al. 2011).

For *P. mira*, PacBio SMRT reads were *de novo* assembled using FALCON (https://github.com/PacificBiosciences/FALCON/). Approximately 13.93 Gb of PacBio SMRT reads were first pairwise compared, and the longest 60 coverage of subreads were selected as seeds to do error correction with parameters ’--output_multi --min_idt 0.70 --min_cov 4 --max_n_read 300 ’. The corrected reads were then aligned to each other to construct string graphs with parameters ‘--length_cutoff_pr 11000’. The graphs were further filtered with parameters ’--max_diff 70 --max_cov 70 --min_cov 3 ’ and contigs were finally generated according to these graphs. All PacBio SMRT reads were mapped back to the assembled contigs with Blast and the Arrow program implemented in SMRT Link (PacBio) was used for error correction with default parameters. The Illumina paired-end reads were then mapped to the corrected contigs to perform the second round of error correction. To further improve the continuity of the assembly, SSPACE (v3.0) was used to build scaffolds using reads from all the mate pair libraries. FragScaff v1-1 (Adey et al., 2014) was further applied to build superscaffolds using the barcoded sequencing reads. Finally, Hi-C data were used to correct superscaffolds and cluster the scaffolds into pseudochromosomes.

To evaluate the quality of the genome assemblies, we first performed BUSCO v3.0.2b (Simao et al., 2015) analysis on the four assembled genomes with the 1,440 conserved plant single-copy orthologs. We then evaluated the assemblies by aligning the RNA-Seq reads to the corresponding assemblies.

### Repetitive element identification

A combined strategy based on homology alignment and *de novo* search was used to identify repeat elements in the four wild peach genomes. For *de novo* prediction of transposable elements (TEs), we used RepeatModeler (http://www.repeatmasker.org/RepeatModeler.html), RepeatScout (http://www.repeatmasker.org/), Piler (Edgar, et al., 2005), and LTR-Finder (Xu et al., 2007) with default parameters. For alignment of homologous sequences to identify repeats in the assembled genomes, we used RepeatProteinMask and RepeatMasker (http://www.repeatmasker.org) with the repbase library (Jurka et al., 2005). Transposable elements overlapping with the same type of repeats were integrated, while those with low scores were removed if they overlapped more than 80 percent of their lengths and belonged to different types.

### Gene prediction and functional annotation

Gene prediction was performed using a combination of homology, *ab initio* and transcriptome based approaches. For homology-based prediction, protein sequences from *P. persica, Pyrus bretschneideri, P. mume, Malus domestica*, and *Fragaria vesca* (Genome Database for Rosaceae; https://www.rosaceae.org) and *Vitis vinifera* (http://www.genoscope.cns.fr/externe/GenomeBrowser/Vitis/) *and Arabidopsis thaliana* (https://www.arabidopsis.org) were downloaded and aligned to the peach assemblies. Augustus (Stanke et al., 2004), GlimmerHMM (Majoros et al., 2004) and SNAP (Korf, I. 2004) were used for *ab initio* predictions. For transcriptome-based prediction, RNA-Seq data derived from root, phloem, leaf, flower, and fruit were mapped to the assemblies using HISAT2 software (Kim et al., 2019) and assembled into the transcripts using Cufflinks (version 2.1.1) with a reference-guided approach (Trapnell et al., 2010). Moreover, RNA-Seq data were also *de novo* assembled using Trinity v2.0 (Grabherr et al., 2011) and open reading frames in the assembled transcripts were predicted using PASA (Haas et al., 2008). Finally, gene models generated from all three approaches were integrated using EvidenceModeler (Haas et al., 2008) (EVM) to generate the final consensus gene models.

The predicted genes were functionally annotated by comparing their protein sequences against the NCBI non-redundant (nr), Swiss-Prot (http://www.uniprot.org/), TrEMBL (http://www.uniprot.org/), Kyoto Encyclopedia of Genes and Genomes (KEGG, http://www.genome.jp/kegg/), InterPro, and GO databases.

tRNAscan-SE (Lowe and Eddy, 1997) was used with default parameters to identify tRNA sequences in the genome assemblies. rRNAs in the genomes were identified by aligning the reference rRNA sequence of relative species to the assemblies using BLAST with *E*-values <1e-10 and nucleotide sequence identities > 95%. Finally, the INFERNAL v1.1 (http://infernal.janelia.org/) software was used to compare the genome assemblies with the Rfam database (http://rfam.xfam.org/) to predict miRNA and snRNA sequences.

### Genome alignment and collinearity analysis

Orthologous genes within the *P. mira* and *P. persica* genomes were identified using BLASTP (E value < 1e^−5^), and MCScanX (Wang et al., 2012b) was used to identify syntenic blocks between the two genomes. The collinearity of the two genomes were then plotted according to the identified synteic blocks.

Four wild peach genomes were aligned to the *P. persica* genome using LASTZ (Harris et al., 2007) with the parameters of ‘M=254K=4500 L=3000 Y=15000 --seed=match 12 --step=20 --identity=85’ (Shi et al., 2017). In order to avoid the interference caused by repetitive sequences in alignments, RepeatMasker and RepBase library were used to mask the repetitive sequences in genomes of *P. persica* and four wild species. The raw alignments were combined into larger blocks using the ChainNet algorithm implemented in LASTZ.

### Variation identification

We identified SNPs and small indels (< 50 bp) between the four wild and reference peach genomes using SAMtools (http://samtools.sourceforge.net/) and LAST (http://last.cbrc.jp) with parameters ‘-m20 -E0.05’, and SNP and indel filtering criteria ‘minimum quality = 20, minimum depth = 5, maximum depth = 200’. SVs were identified from genome alignments by LAST with parameters‘-m20 -E0.05’. CNVs were identified using CNVnator-0.3.3 (Abyzov et al., 2011).

### Promoter activity measurement

A total of five primers upstream and one downstream of the start codon of *Prupe.2G053600* were synthesized and used to amplify a series of 5 indel regions in the *Prupe.2G053600* promoter using PCR amplification. The amplified PCR products were ligated into pGEM-T easy vector and cloned into pBI101 binary vector after digested by XbaI and BamHI. Furthermore, each of the 5 amplified products was transformed into *Agrobacterium tunefaciens* (GV1301) cells and collected and resuspended in infiltration buffer, and then transformed into 6-week-old tobacco leaves using sterilized syringes. The transiently transformed tobacco plants were grown in a growth chamber for 48 h and the infection sites were cut to measure glucurinidase (GUS) activity as described in Jafferson et al. (1987). The pBI121 vector was used as a positive control.

### Transgenic analysis

The full-length open reading frame of the *Prupe.2G053600* gene was amplified through PCR using cDNA synthesized from RNA that was isolated from root of the ‘Hong Gen Gan Su Tao 1#’ (*P. kansuensis*). The amplified product was cloned into the pEASY vector driven by the cauliflower mosaic virus (CaMV) 35 S promoter. The resulting vector was transformed into *Solanum lycopersicum* cv. Micro-Tom by *Agrobacterium tumefaciens* C58. The T0 plants were generated and inoculated with *M. incognita* to observe resistance and measure gene expression.

Similarly, one candidate gene, *evm.model.Pm01.401*, was cloned from the leaf of ‘2210-198’ (*P. mira*) and ligated to the vector and transformed into *A. thaliana* ‘Columbia’. When the transformed plants were grown to about 5 leaves, low temperature (0 °C) treatment was applied and samples were collected at 0, 24, and 48 h post treatment. Phenotype was observed and the drooping leaves were used to indicate that the accession was susceptible to low temperature.

### Comparative analysis

Protein sequences from 11 plant species including *P. persica* (phytozomev10), *P. mira, P. davidiana, P. kansuensis, P. ferganensis, Prunus dulcis* (https://www.rosaceae.org/species/prunus/prunus_dulsis/lauranne/genome_v1.0), *Prunus tangutica* (derived from transcripts *de novo* assembled from RNA-Seq data), *Prunus mume* (http://prunusmumegenome.bjfu.edu.cn/index.jsp), *Fragaria vesca* (phytozome v10), *A. thaliana* (phytozome v10), and *Vitis vinifera* (phytozome v10) were used to construct orthologous gene families. To remove redundancy caused by alternative splicing, we retained only the gene model at each gene locus that encoded the longest protein. To exclude putative fragmented genes, genes encoding protein sequences shorter than 50 amino acids were filtered out. All-against-all BLASTp was performed for these protein sequences with an E-value cut-off of 1e-5. OrthoMCL V1.4 (Li et al., 2003) was then used to cluster genes into gene families with the parameter ‘-inflation 1.5’.

Protein sequences from 3,548 single-copy gene families were used for phylogenetic tree construction. MUSCLE (Edgar et al., 2004) was used for multiple sequence alignment for protein sequences in each single-copy family with default parameters. The alignments from all single-copy families were then concatenated into a super alignment matrix, which was used for phylogenetic tree construction using the Maximum likelihood (ML) method implemented in the PhyML software (http://www.atgc-montpellier.fr/phyml/binaries.php). Divergence times between the 11 species were estimated using MCMCTree in PAML software (http://abacus.gene.ucl.ac.uk/software/paml.html) with the options ‘correlated rates’ and ‘JC69’ model. A Markov Chain Monte Carlo analysis was run for 10,000 generations, using a burn-in of 10,000 iterations and sample-frequency of 2. Three calibration points were applied according to the TimeTree database (http://www.timetree.org): *A. thaliana* and *V. vinifera* (103.2-119.5 Mya), *A. thaliana* and the common ancestor of *M. domestica, P. mume*, and *P. persica* (97.1-109.0 Mya), *P. mume* and other *Prunus* species (17.1-34.0 Mya).

To detect the whole genome duplication events, we first identified collinearity blocks using paralogous gene pairs with software MCScanX (Wang et al., 2012b). Using the sum of transversion of fourfold degenerate site divided by the sum of fourfold degenerate sites, we then calculated 4dTv (transversion of fourfold degenerate site) values of each block. In addition, Ks values of homologous gene pairs were also calculated using PAML (Yang, 2007) based on the sequence alignments by MUSCLE (Edgar, 2004), to validate speciation times.

### Gene family expansions and contractions

Expansion and contractions of orthologous gene families were determined using CAFE (De Bie et al., 2006), which uses a birth and death process to model gene gain and loss over a phylogeny. Significance of changes in gene family size in a phylogeny was tested by calculating the p-value on each branch using the Viterbi method with a randomly generated likelihood distribution. This method calculates exact p-values for transitions between the parent and child family sizes for all branches of the phylogenetic tree. Enrichment of GO terms and KEGG pathways in the expanded gene families of each of the four wild peach species were identified using the R package clusterProfiler (Yu et al., 2012).

### Identification of nonlinear evolution event of *P. davidiana*

Using SNPs identified from 126 peach accessions, we constructed a neighbor joining tree with 1000 bootstraps using TreeBeST 1.9.2 (Vilella et al., 2009). We then investigated population structure using the program frappe (Tang et al., 2005) with the number of assumed genetic clusters (K) ranging from two to five, and 10 000 iterations for each run. We also performed PCA to evaluate the evolution path using the software GCTA (Yang et al., 2011).

To trace the origin of genes in *P. davidiana*, each gene was aligned to other genomes to calculate the alignment score. Genes were classified as putatively originating from the specie which had the highest alignment scores. Finally, we clustered all contigs of *P. davidiana* into pseudomolecules using the Hi-C technology. Simultaneously, collinearity among *P. dulcis, P. davidiana*, and *P. mira* was plotted based on the identified syntenic blocks.

### Resistance genes

Hidden Markov model search (HMMER; http://hmmer.janelia.org) was used to identify R genes in the four wild peach genomes according to the NBS (NB-ARC) domain (PF00931), TIR model (PF01582), and several LRR models (PF00560, PF07723, PF07725, PF12799, PF13306, PF13516, PF13504, PF13855 and PF08263) in the Pfam database (http://pfam.sanger.ac.uk). CC motifs were detected using the COILS prediction program 2.2 (https://embnet.vital-it.ch/software/COILS_form.html) with a p score cut-off of 0.9.

### Identification of selective sweeps associated with high-altitude adaptation in *P. mira*

Raw genome reads of the 32 accessions of *P. mira* from Tibet, China with different altitudes were processed to remove adaptor, contaminated and low quality sequences, and the cleaned reads were mapped to the assembled *P. mira* genome using BWA (version 0.7.8) (Li and Durbin, 2009). Based on the alignments, the potential PCR duplicates were removed using the SAMtools command “rmdup”. SNP calling at the population level was performed using SAMtools (Li et al., 2009). The identified SNPs supported by at least five mapped reads, mapping quality ≥20, and Phred-scaled genotype quality ≥5, and with less than 0.2 missing data were considered high-quality SNPs (1,394,483), and used for subsequent analyses.

We first constructed a phylogenetic tree and performed structure analysis of the 32 accessions of *P. mira* to remove accessions with admixture background. To identify genome-wide selective sweeps associated with high-altitude adaptation, we scanned the genome in 50-kb sliding windows with a step size of 10 kb, and calculated the reduction in nucleotide diversity (π) based on the *P. mira* accessions originating in high-altitude to low-altitude regions (π_high_/π_low_). In addition, selection statistics (Tajima’s D) and population differentiation (*F*_*ST*_) between the two groups were also calculated. Windows with the top 5% of the π ratios, Tajima’s D ratio and *F*_ST_ values were considered as selective sweeps.

## DATA AVAILABILITY

Raw resequencing data for 108 of 158 peach accessions generated in this study have been deposited into the NCBI database as a BioProject under accession PRJNA645279, and for the other 40 have been deposited previously under accession numbers SRP168153 and SRP173101. The assemblies of four genomes have been uploaded to Genome Database for Rosaceae (https://www.rosaceae.org).

## ACKNOWLEDGMENTS

This study was supported by grants from the National Key Research and Development Program (2019YFD1000203), the Agricultural Science and Technology Innovation Program (CAAS-ASTIP-2019-ZFRI-01), and National Horticulture Germplasm Resources Center.

## AUTHOR CONTRIBUTIONS

L.W. and K.C. conceived the project. Y.L. and X.Z. contributed to the original concept of the project. G.Z., W.F., C.C., X.W., and J.W. collected samples and performed phenotyping. K.L. conducted gene expression analysis and transgenic experiments. K.C., S.D., Z.P., and Z.F. analysed the data. K.C. wrote the paper.

## COMPETING FINANCIAL INTERESTS

The authors declare no competing financial interests.

## ETHICS APPROVAL AND CONSENT TO PARTICIPATE

Not applicable.

## Additional files

**Additional file 1**

Supplementary Table 1 List of 158 peach samples and summary statistics of genome resequencing.

Supplementary Table 2 Assembling statistics of reads that were not mapped to the peach reference genome.

Supplementary Table 3 Summary statistics of *P. persica* pan-genome.

Supplementary Table 4 Functional annotations of *P. persica* non-reference genes.

Supplementary Table 5 Expression of 138 of 923 non-reference (novel) genes which expressed at >1 reads per kilobase of exon per million mapped reads (RPKM) in at least 1 of 5 tissues in *P. persica*.

Supplementary Table 6 Validation of non-reference sequences using the Sanger technology.

Supplementary Table 7 Genome survey of four wild peach species (kmer = 17).

Supplementary Table 8 Summary of genome sequencing of four wild peach species.

Supplementary Table 9 Pseudochromosome lengths of the *P. mira* assembly.

Supplementary Table 10 BUSCO analysis of the genome assemblies of four wild peach species.

Supplementary Table 11 Mapping statistics of RNA-Seq reads to the corresponding genome assemblies of four wild peach species.

Supplementary Table 12 Statistic of repeat sequences in the assemblies of four wild peach species.

Supplementary Table 13 Prediction of protein-coding genes in the genomes of four wild peach species.

Supplementary Table 14 Statistics of predicted protein-coding genes in four wild peach species compared to other species.

Supplementary Table 15 Statistics of gene functional annotation in the four wild peach species.

Supplementary Table 16 Non-coding RNAs identified in genomes of four wild wild peach species.

Supplementary Table 17 SNPs identified between genomes of each of the four wild species and *P. persica*.

Supplementary Table 18 Small indels (<50 bp) identified between genomes of each of the four wild species and *P. persica*.

Supplementary Table 19 Genome structural variants (≥ 50 bp) between the four wild species and *P.persica*.

Supplementary Table 20 Statistics of copy number variations between the four wild species and *P. persica*.

Supplementary Table 21 Variations in the promoter and mRNA regions of *R* genes on Chr. 2 (5-7 Mb) that were specific to *P. kansuensis*.

Supplementary Table 22 Statistics of resistance genes in the four wild peach species.

Supplementary Table 23 Genes selected between the two subgroups of *P. mira* which originated from high- and low-altitude regions.

Supplementary Table 24 Selected genes in the KEGG pathways associated with plateau adaptability.

**Additional file 2**

Supplementary Figure 1 Distribution of genes from the pan-genome of *P. persica* in different populations.

Supplementary Figure 2 Linalool contents of mature fruits in 57 peach varieties evaluated in 2015 and 2016.

Supplementary Figure 3 Expression of reference and non-reference genes in different tissues.

Supplementary Figure 4 Sequence alignment of PCR products and novel non-reference sequences.

Supplementary Figure 5 Pictures of the *P. mira* accession 2010-138 used for genome assembly. (a) Sampling location (red dot) shown on the map. (b) Tree of *P. mira* accession 2010-138.

Supplementary Figure 6 Estimation of genome sizes of *P. mira* (a), *P. davidiana* (b), *P. kansuensis* (c), and *P. ferganensis* (d) based on K-mer analysis.

Supplementary Figure 7 Characteristics of the *P. mira* and *P. persica* genomes. The outermost to innermost tracks indicate repeat sequence density (a), gene density (b), gene expression in fruit (c), flower (d), leaf (e), and seed (f), and GC content (g) of *P. mira* (Pm) and *P. persica* (Pp). Lines in the center of the circle indicate syntenic regions between the different chromosomes of *P. mira* and *P. persica*.

Supplementary Figure 8 Genome variations across the pseudo-chromosomes of four wild peach species compared to the reference (*Prunus persica*). The circles from the outer to the inner (A-P) represent copy number variation (CNV) density in *P. ferganensis* (A), *P. kansuensis* (B), *P. davidiana* (C), and *P. mira* (D), and structure variations (SVs) in *P. ferganensis* (E), *P. kansuensis* (F), *P. davidiana* (G), and *P. mira* (H), and indels in *P. ferganensis* (I), *P. kansuensis* (J), *P. davidiana* (K), and *P. mira* (L), as well as SNPs in *P. ferganensis* (M), *P. kansuensis* (N), *P. davidiana* (O), and *P. mira* (P) in each sliding window of 0.1 Mb.

Supplementary Figure 9 KEGG pathways enriched in genes comprising large-effect SNPs of between *P. ferganensis* and *P. persica*.

Supplementary Figure 10 KEGG pathways enriched in genes comprising indels in four wild peach species compared to *P. persica*.

Supplementary Figure 11 KEGG pathways enriched in genes comprising structure variations in *P. mira and P. kansuensiss* compared to *P. persica*.

Supplementary Figure 12 KEGG pathways enriched in genes comprising duplication (a-d) and deletion (e-h) copy number variations in *P. mira* (a, e), *P. davidiana* (b, f). *P. kansuensis* (c, g), and *P. ferganensis* (d, h) compared to *P. persica*.

Supplementary Figure 13 Venn diagram of gene families identified from the five species of peach (a) and expanded (b), and contracted (c) gene families from the four wild species compared to *P. persica*.

Supplementary Figure 14 Statistics of single-copy orthologs, multiple-copy orthologs, and unique orthologs in 11 species.

Supplementary Figure 15 Whole-genome duplication and speciation events in peach as revealed by the distribution of 4DTv distance among paralogous and orthologs genes in different species.

Supplementary Figure 16 Percent of *P. davidiana*-specific contigs covered by reads from different *Prunus* species.

Supplementary Figure 17 Geographical distribution of *P. mira* (red circle), *P. davidiana* (yellow), and *P. dulcis* (orange) which originated in China.

Supplementary Figure 18 Distribution of resistance (*R*) genes across the 8 chromosomes in five peach species of peach and their overlaps with disease resistance QTLs.

Supplementary Figure 19 KEGG pathways enriched in expanded and contracted gene families of the four wild peach species.

Supplementary Figure 20 Phylogenetic tree of 32 accessions of *P. mira* (a) originating from regions with different altitudes (b) and the population structure (c) when K=2 and 3.

Supplementary Figure 21 Two genome regions associated with high-altitude adaptation.

## Notes

### Competing Interest Statement

The authors have declared no competing interest.

