## Supplementary figure 1-21 for "Pan-genome analyses of peach and its wild relatives provide insights into the genetics of disease resistance and species adaptation"

**Supplementary Figure 1 Distribution of genes from the pan-genome of *P. persica* in different populations.**

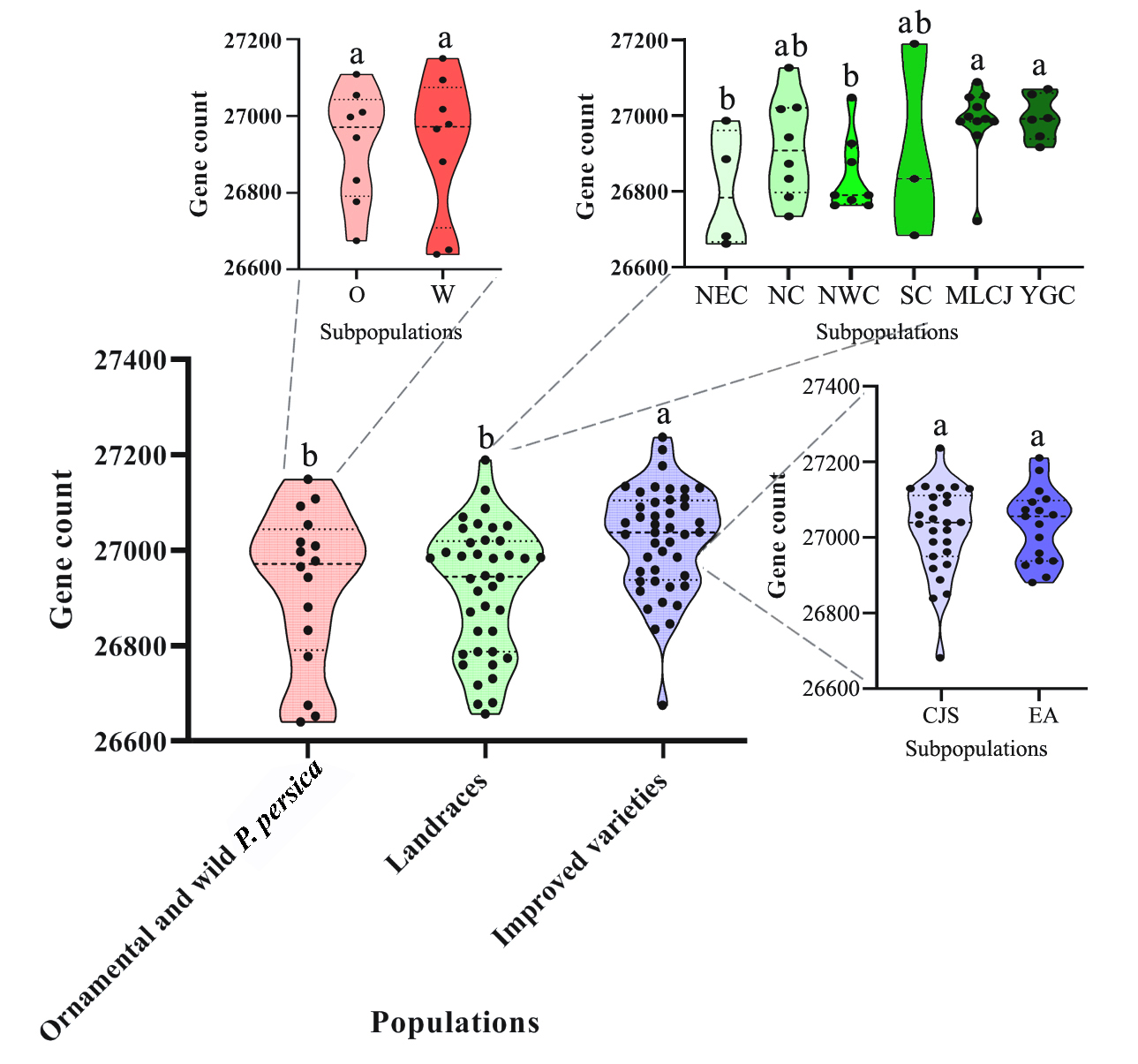

‘O’ indicated the ornamental peaches. ‘W’ indicated wild peaches belonging *P. persica* that can be used as rootstock of improved varieties. And among of landraces, the accessions were divided into six different geographical populations, such as northwest China (NWC), the YunGui plateau (YGC), northeast China (NEC), northern China (NC), the middle and lower reaches of the Changjiang River (MLCJ), and southern China (SC). Among of improved varieties, the accessions were divided into China, Japan and South Korea (CJS), Europe and America (EA) according to their breeding location. And ‘a’ and ‘b’ indicate statistical significance. Observing the distribution of these genes in different accessions, we found genomes of ornamental and wild accessions as well as landraces encoded significantly fewer genes than improved varieties, suggesting a general trend of gene gain during peach improvement and contradictory with the results on the tomato (Gao et al., 2019). Furthermore, more genes were found in the middle and lower reaches of the Changjiang River or YunGui plateau than northwest China or northeast China populations among different geographical populations, might relate to the complexity of their originate background.

**Supplementary Figure 2 Linalool contents of mature fruits in 57 peach varieties evaluated in 2015 and 2016.**

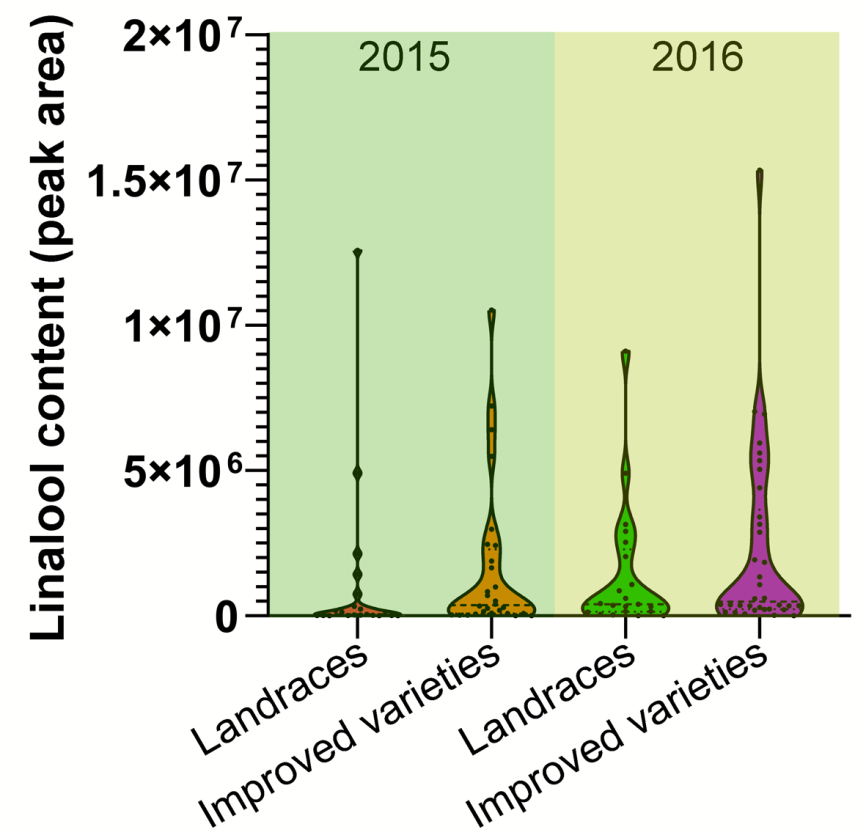

**Supplementary Figure 3 Expression of reference and non-reference genes in different tissues.**

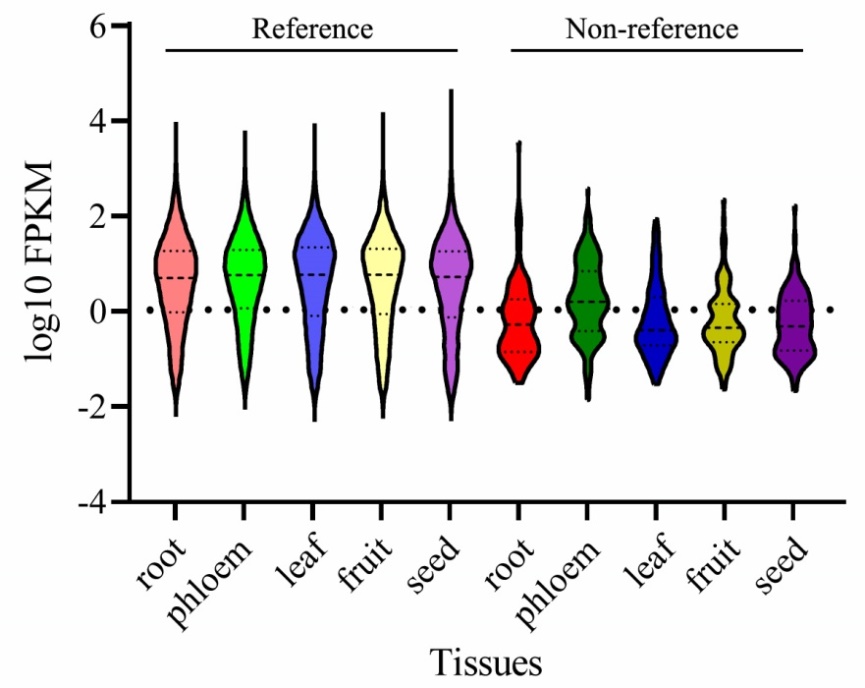

RNA-sequencing (RNA-seq) results indicated that 72.75% of the reference and 14.95% of the non-reference (novel) genes were expressed at >1 reads per kilobase (kb) of exon per million mapped reads (RPKM) in at least 1 of 5 tissues, including young root, phloem, young leaf, fruit (80% ripe) and seed, of a cultivated peach accession, *P. persica* var. Chinese cling. Among them, non-reference genes showed lower expression generally than that of reference ones. Within the non-reference genes, a total of 104 genes showed higher expression in phloem than in the other tissues. Of which, 34 were over-representative in different biological pathways, including 10 genes involving in oxidative phosphorylation and 7 involving in photosynthesis. For example, there were 2 genes encoding photosystem II reaction center protein, 2 genes encoding chlorophyll a-b binding protein, 2 genes encoding ribulose bisphosphate carboxylase/oxygenase activase, and 1 gene encoding photosystem I reaction center protein.

**Supplementary Figure 4 Sequence alignment of PCR products and novel non-reference sequences.**

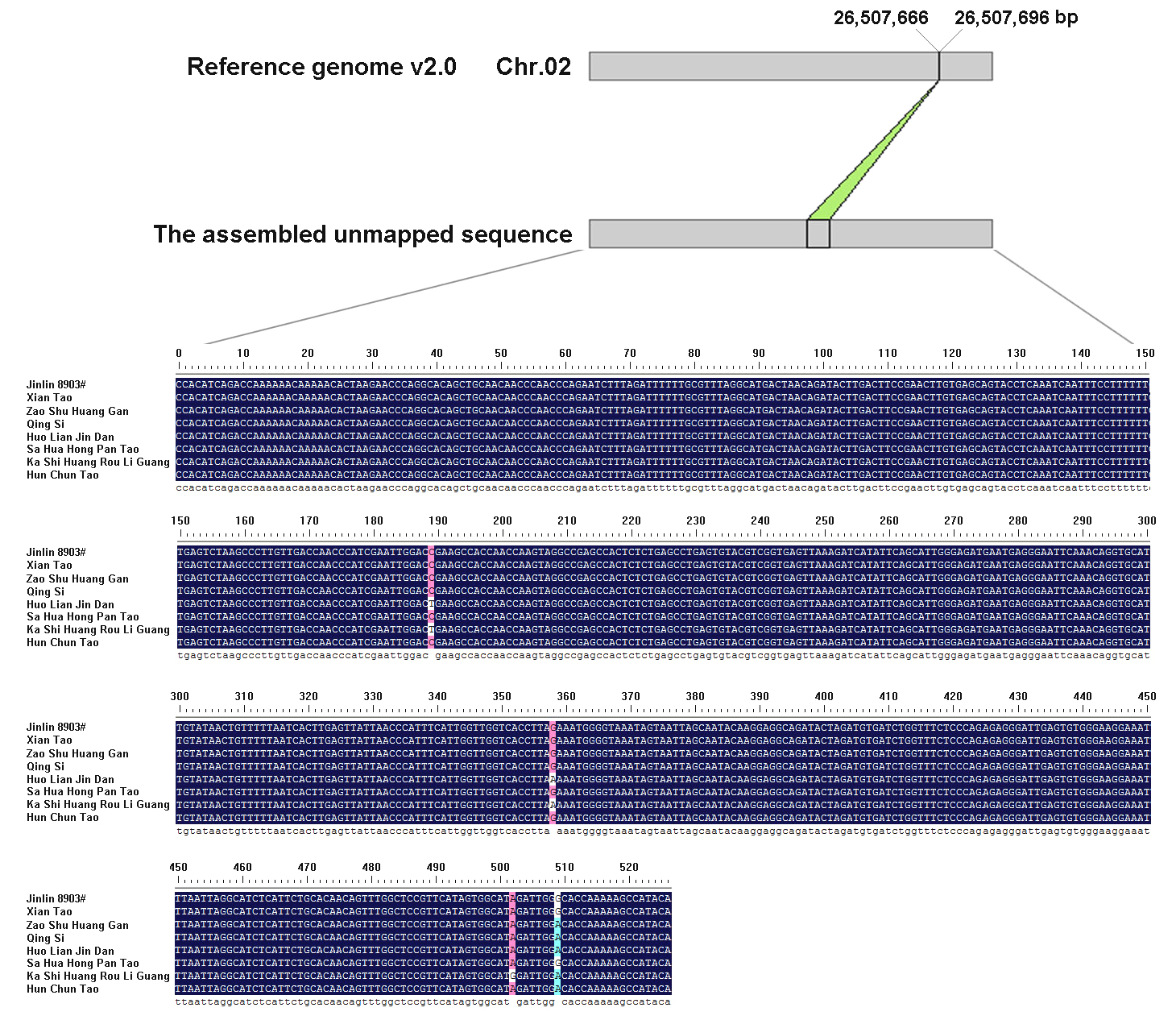

As a result of the average length of these non-reference genes were substantially shorter than that of the whole genome (634 bp versus 2520 bp, Supplementary Table 4), we deduced that positive false might be existed. Therefore, we validated the quality of these assemblers by designing the PCR primers to amplify DNA fragments from the 10 randomly selected contigs in 8 accessions. The Sanger sequencing of these PCR product showed that all of them showed high sequence identity (> 95%) to the assemblers, confirming the high assembled quality of the unmapped sequence of *P. persica* genome. For example, in the above figure, the primers were designed according to one of 10 selected contig, Contig3_A100, and the result indicated the sequence has a low identity with reference genome and the sequence was reliable after compared among different varieties.

**Supplementary Figure 5 Pictures of the *P. mira* accession 2010-138 used for genome assembly. (a) Sampling location (red dot) shown on the map. (b) Tree of *P. mira* accession 2010-138.**

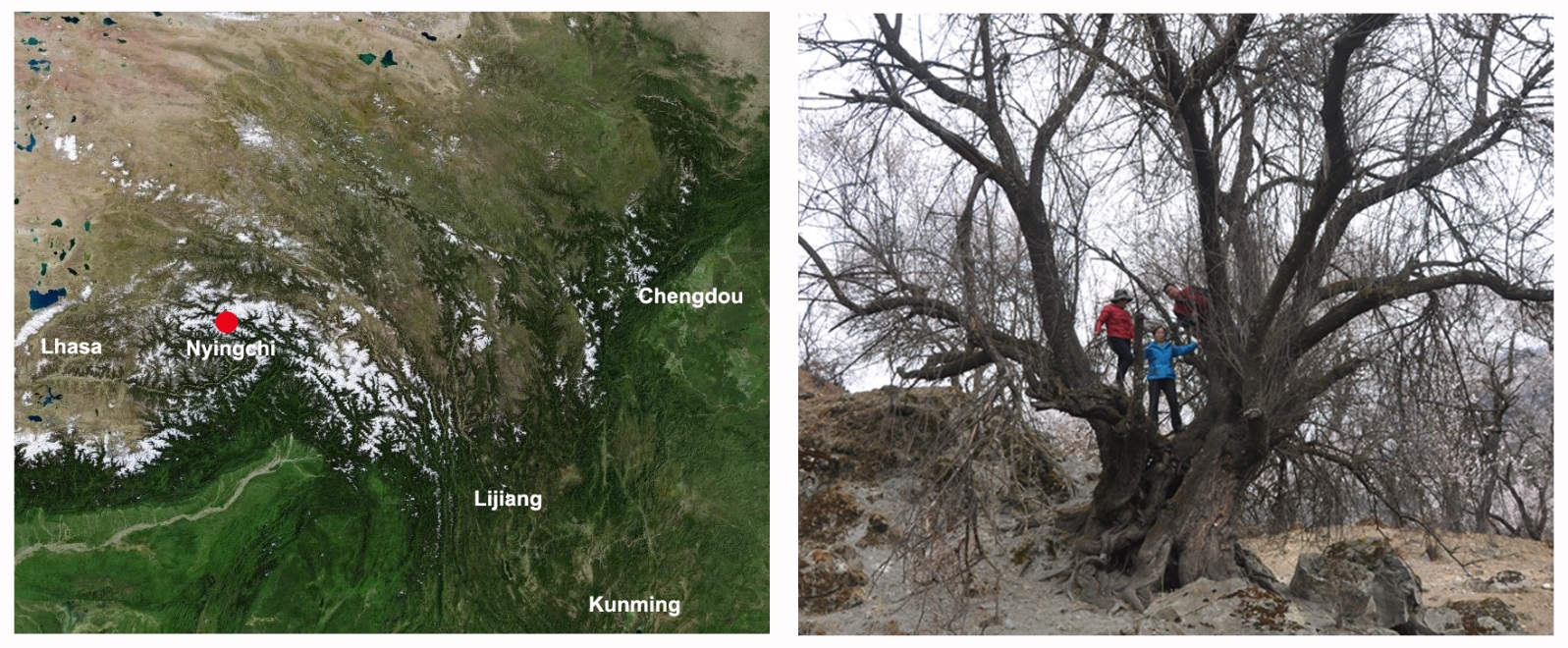

**Supplementary Figure 6 Estimation of genome sizes of *P. mira* (a), *P. davidiana* (b), *P. kansuensis* (c), and *P. ferganensis* (d) based on K-mer analysis.**

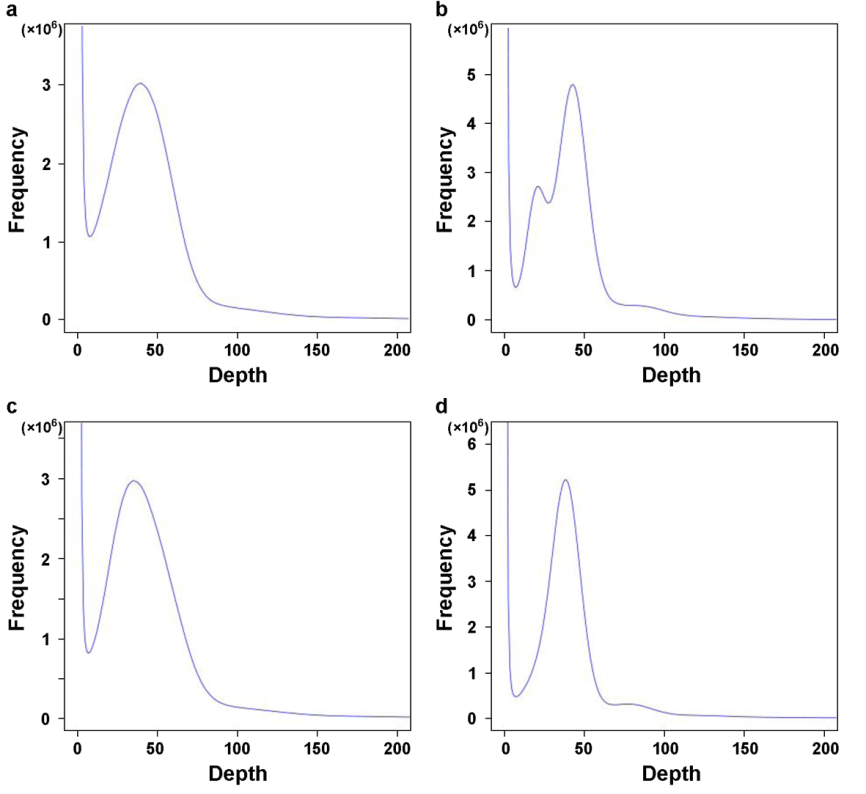

**Supplementary Figure 7 Characteristics of the *P. mira* and *P. persica* genomes.** The outermost to innermost tracks indicate repeat sequence density (a), gene density (b), gene expression in fruit (c), flower (d), leaf (e), and seed (f), and GC content (g) of *P. mira* (Pm) and *P. persica* (Pp). Lines in the center of the circle indicate syntenic regions between the different chromosomes of *P. mira* and *P. persica*.

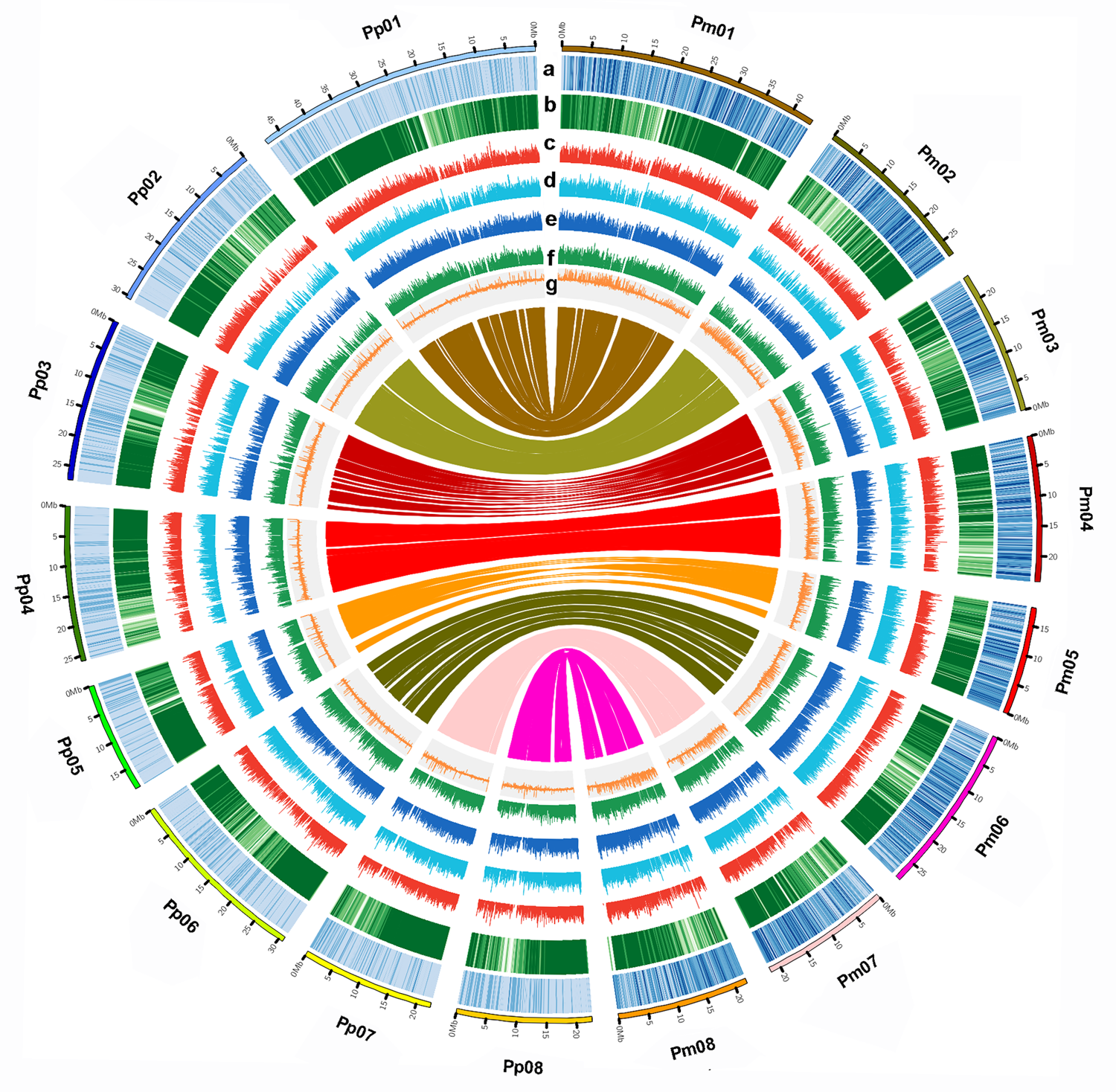

**Supplementary Figure 8 Genome variations across the pseudo-chromosomes of four wild peach species compared to the reference (*Prunus persica*).** The circles from the outer to the inner (A-P) represent copy number variation (CNV) density in *P. ferganensis* (A), *P. kansuensis* (B), *P. davidiana* (C), and *P. mira* (D), and structure variations (SVs) in *P. ferganensis* (E), *P. kansuensis* (F), *P. davidiana* (G), and *P. mira* (H), and indels in *P. ferganensis* (I), *P. kansuensis* (J), *P. davidiana* (K), and *P. mira* (L), as well as SNPs in *P. ferganensis* (M), *P. kansuensis* (N), *P. davidiana* (O), and *P. mira* (P) in each sliding window of 0.1 Mb.

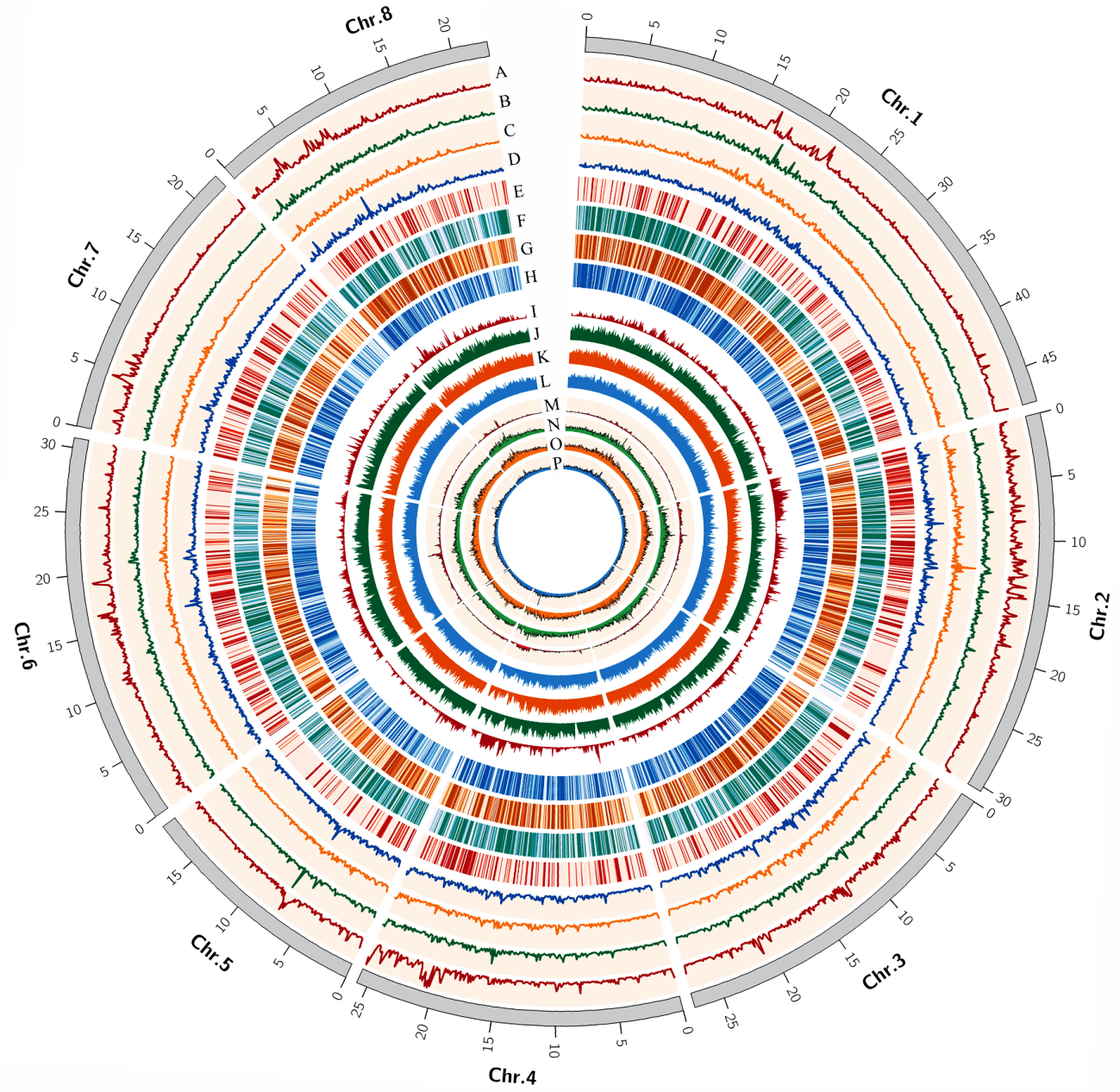

**Supplementary Figure 9 KEGG pathways enriched in genes comprising large-effect SNPs of between *P. ferganensis* and *P. persica*.**

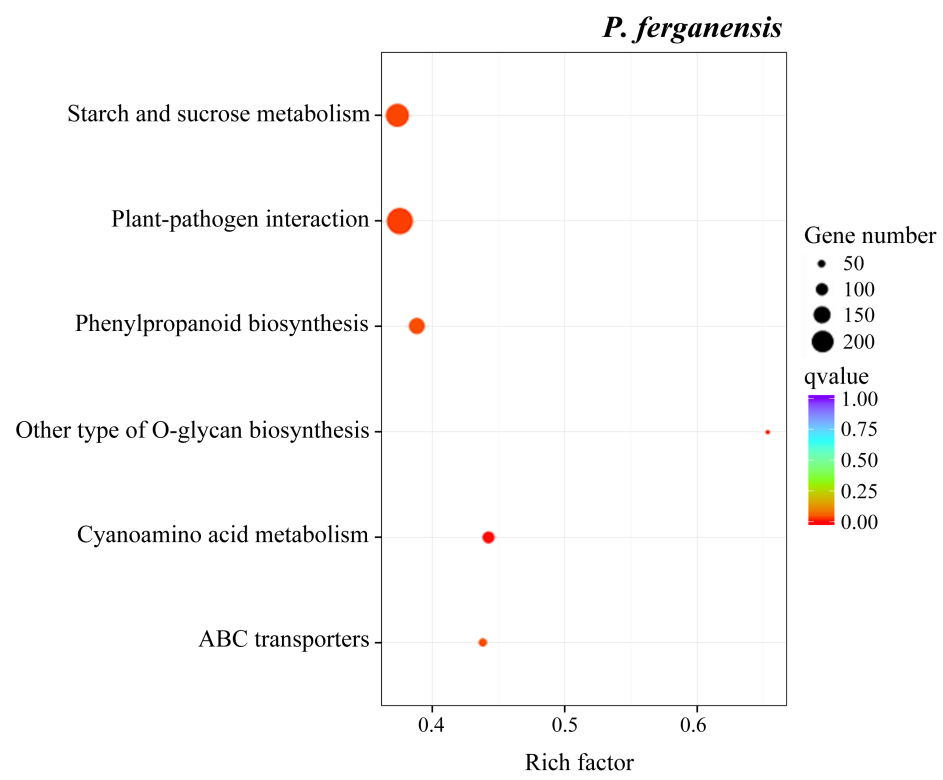

The results show that only *P. ferganensis* has a significant enrichment pathway.

**Supplementary Figure 10 KEGG pathways enriched in genes comprising indels in four wild peach species compared to *P. persica*.**

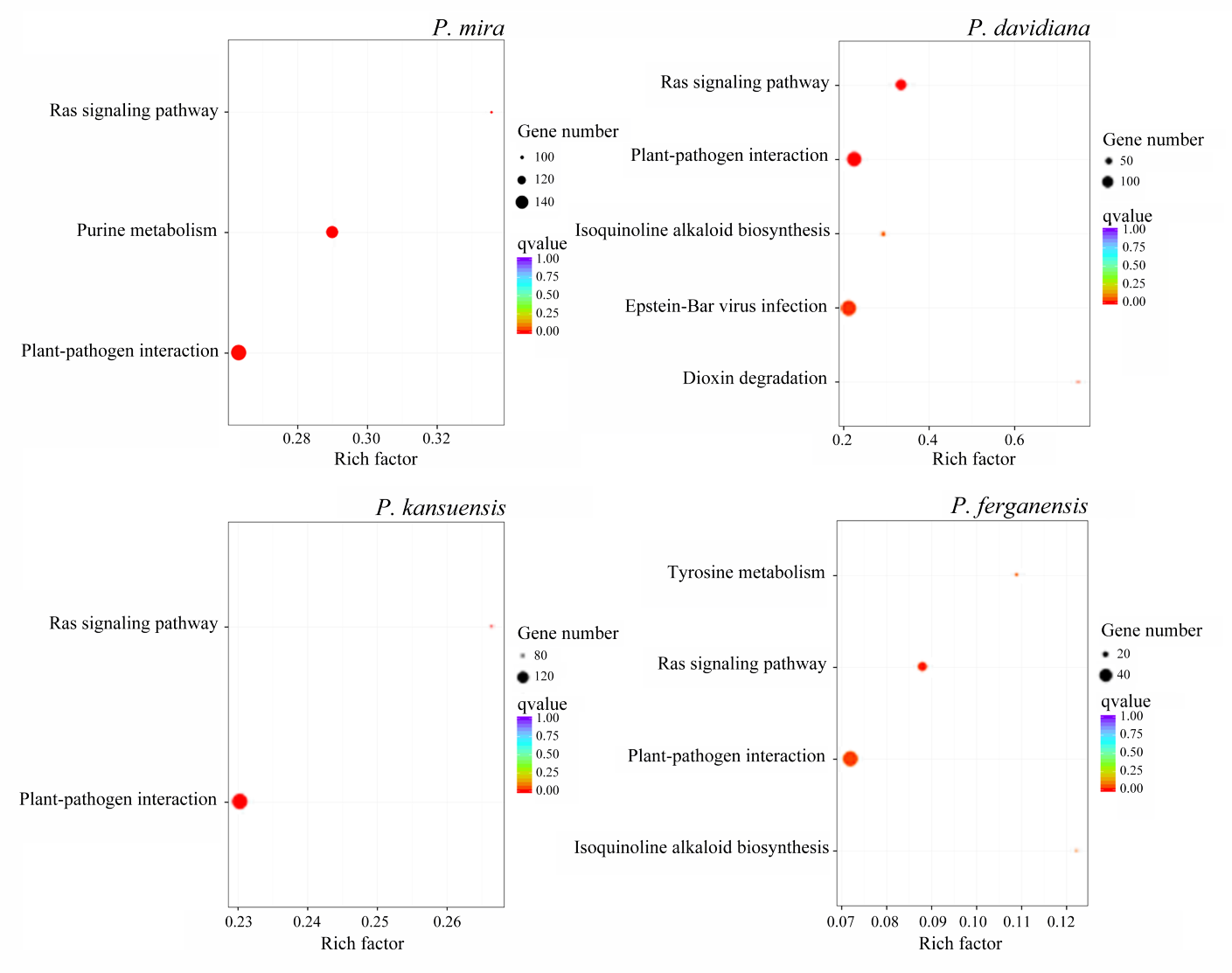

**Supplementary Figure 11 KEGG pathways enriched in genes comprising structure variations in *P. mira and P. kansuensiss* compared to *P. persica*.**

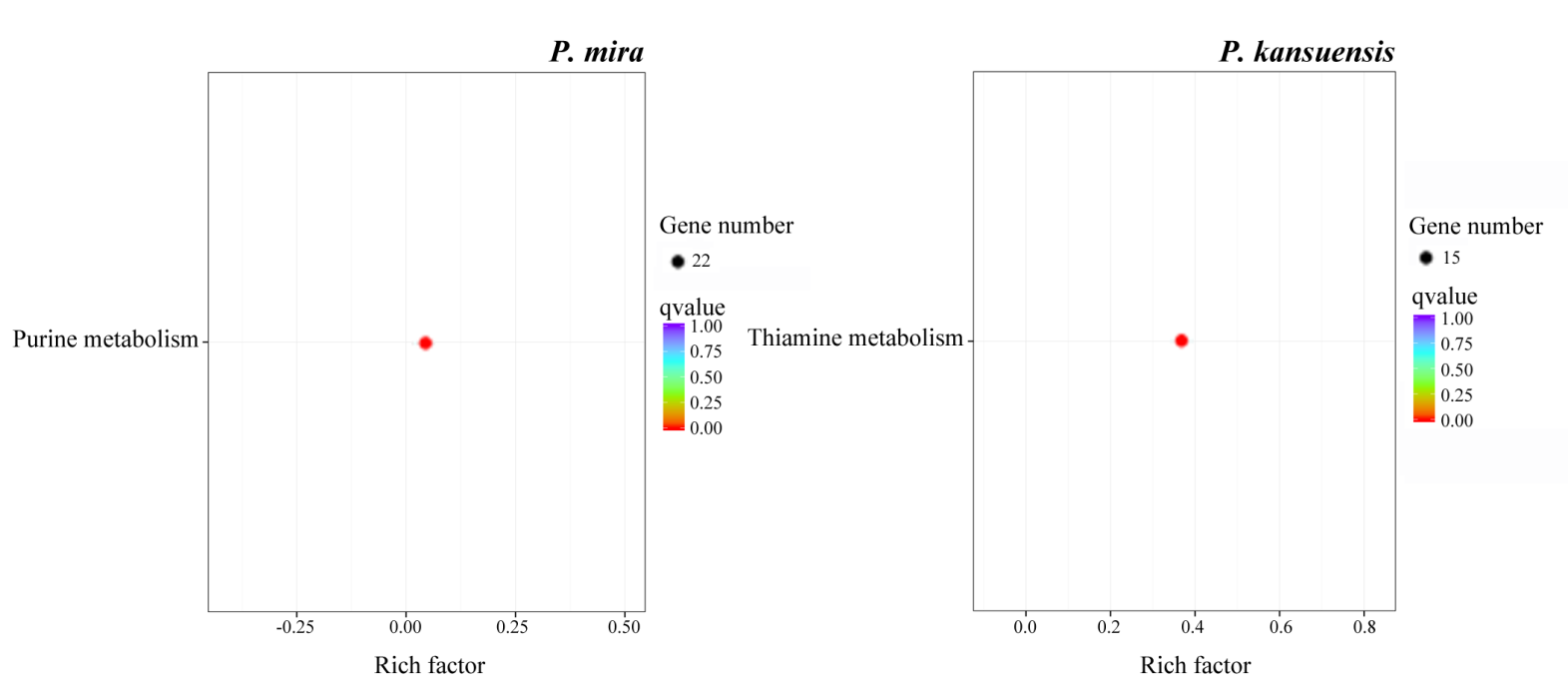

The results show that only *P. mira and P. kansuensiss* have a significant enrichment pathway.

**Supplementary Figure 12 KEGG pathways enriched in genes comprising duplication (a-d) and deletion (e-h) copy number variations in *P. mira* (a, e), *P. davidiana* (b, f). *P. kansuensis* (c, g), and *P. ferganensis* (d, h) compared to *P. persica*.**

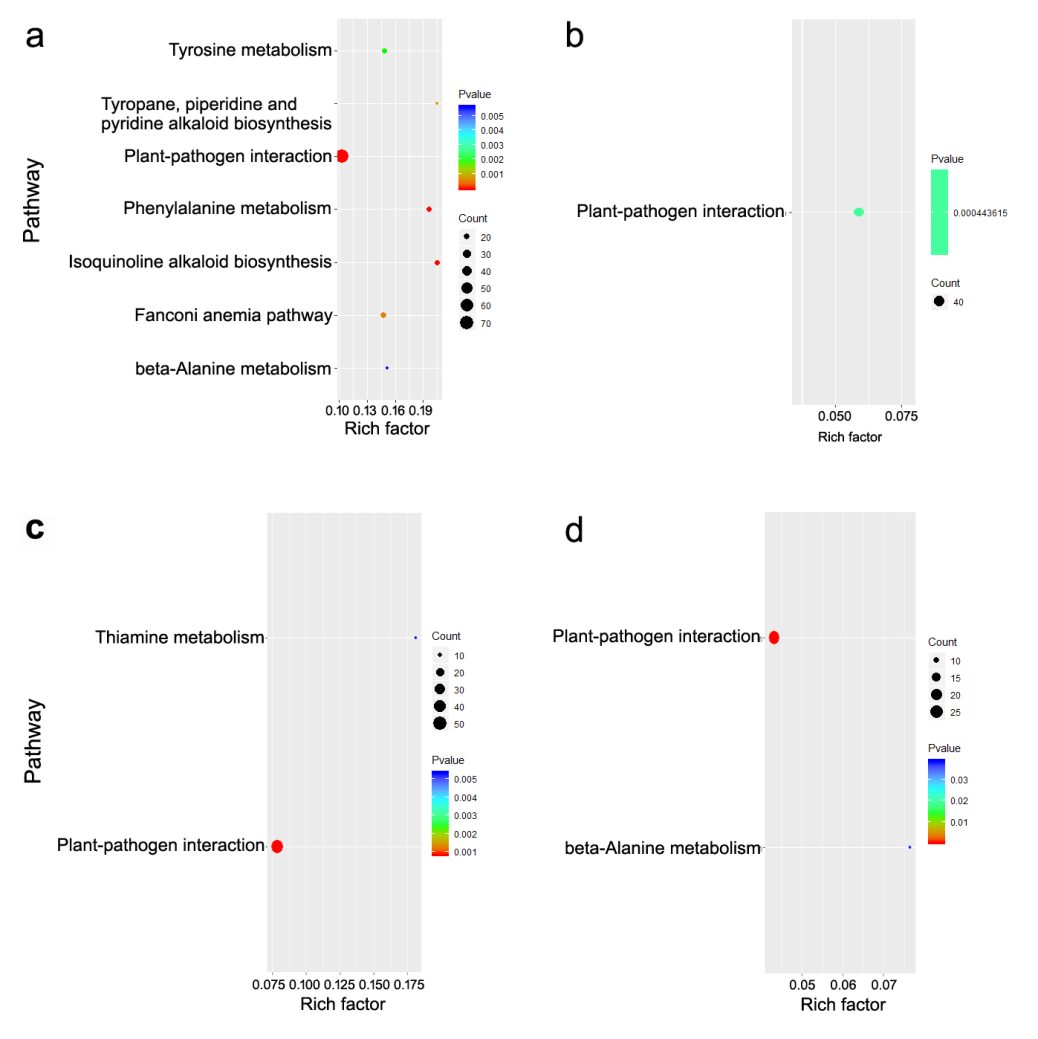

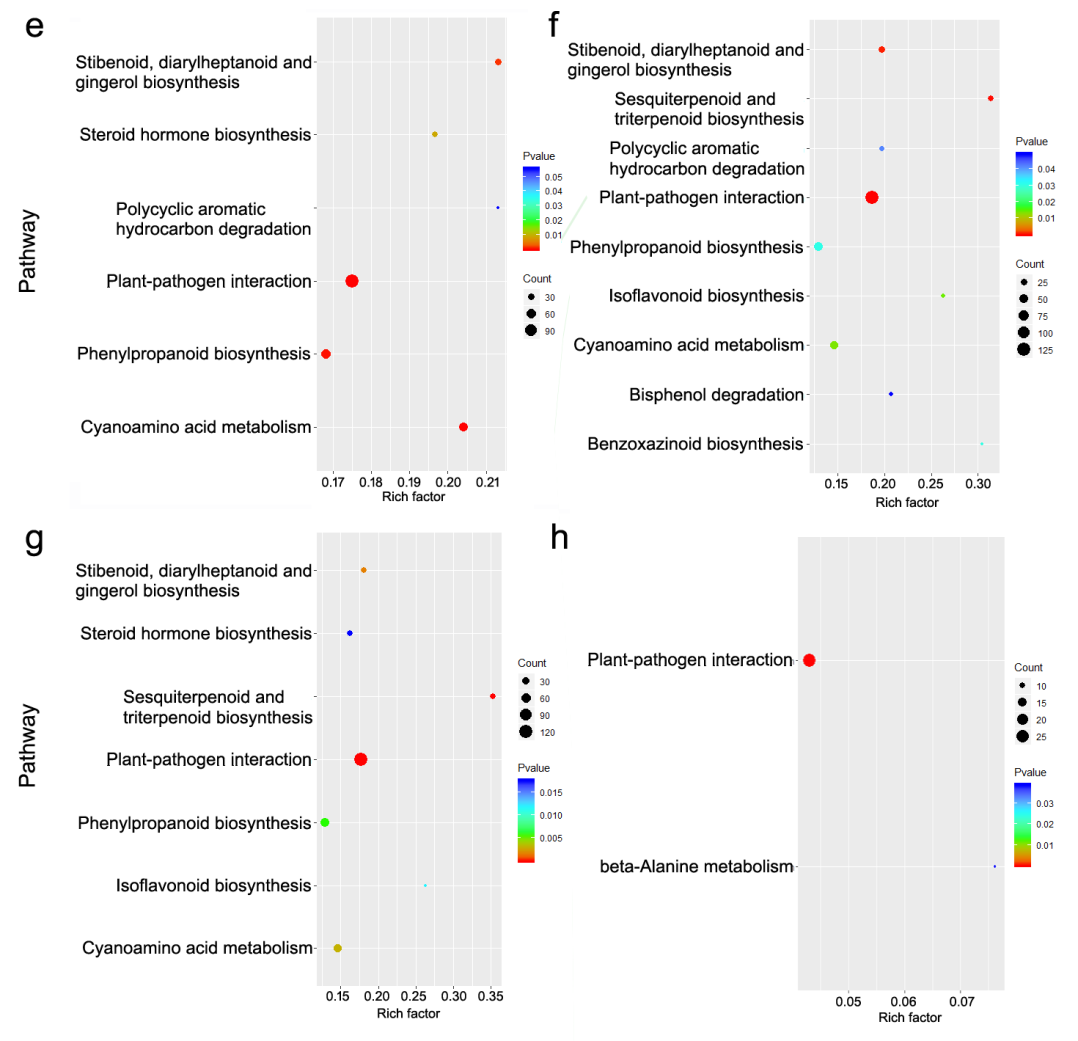

**Supplementary Figure 13 Venn diagram of gene families identified from the five species of peach (a) and expanded (b), and contracted (c) gene families from the four wild species compared to *P. persica*.**

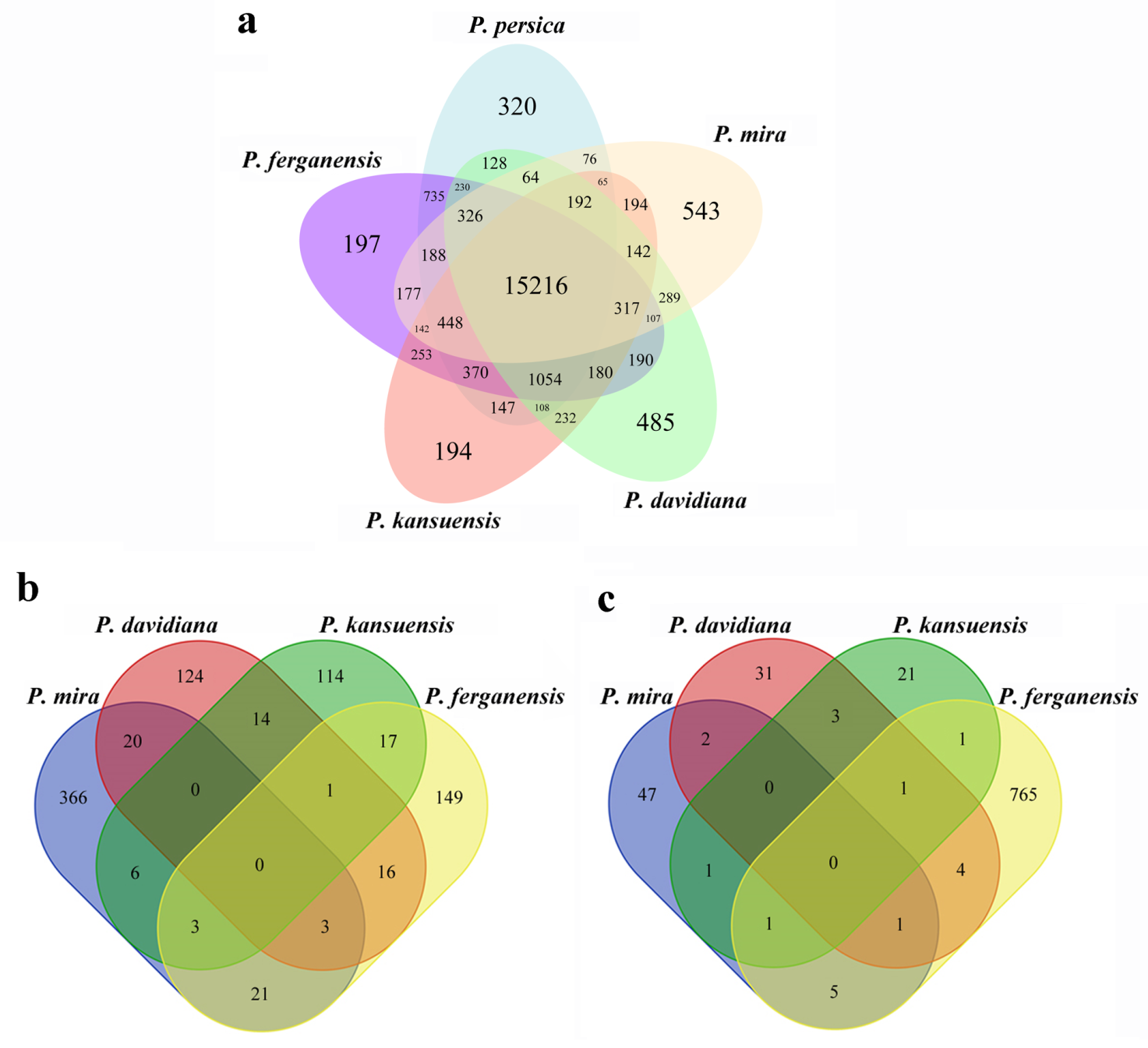

**Supplementary Figure 14 Statistics of single-copy orthologs, multiple-copy orthologs, and unique orthologs in 11 species.**

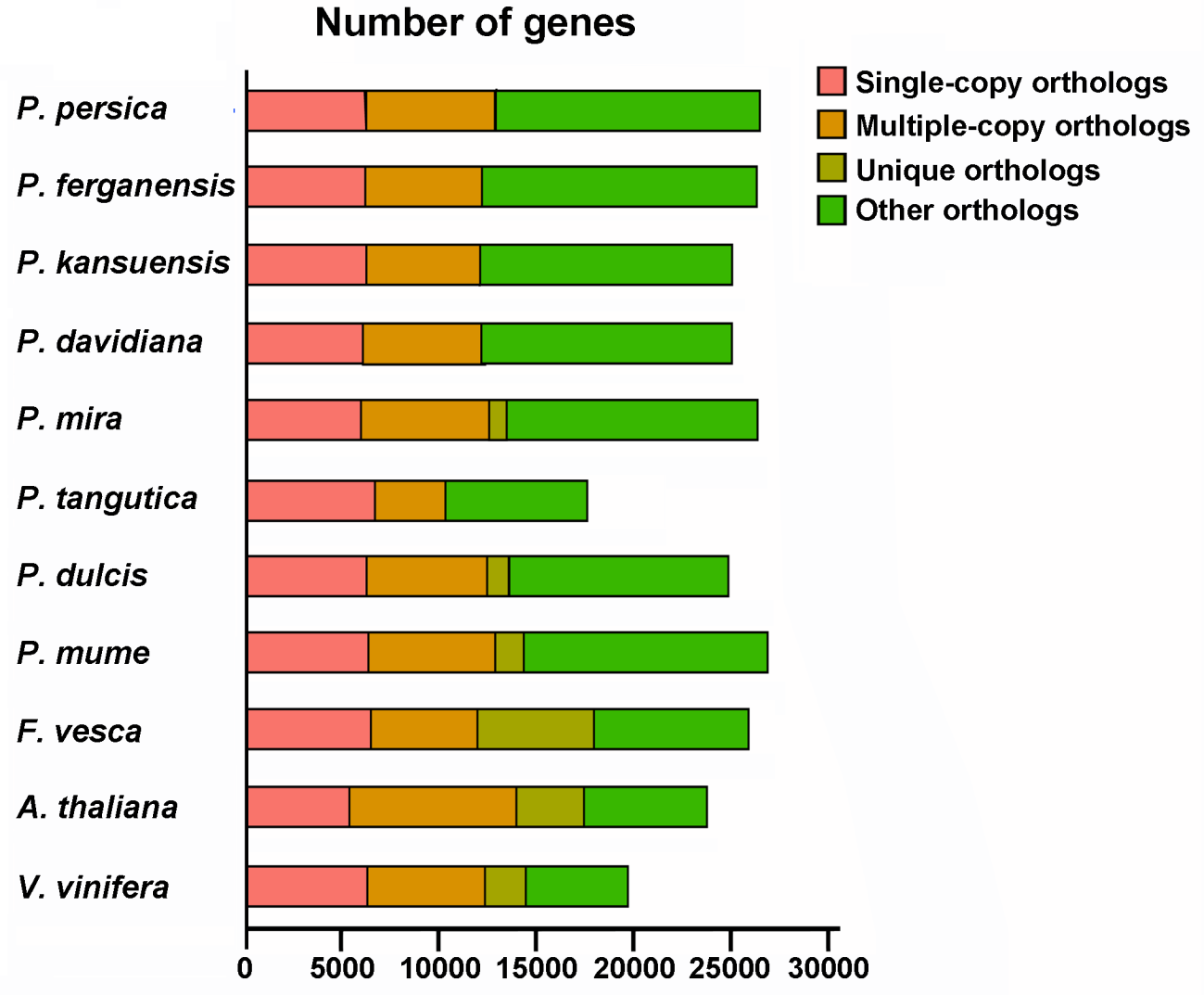

**Supplementary Figure 15 Whole-genome duplication and speciation events in peach as revealed by the distribution of 4DTv distance among paralogous and orthologs genes in different species.**

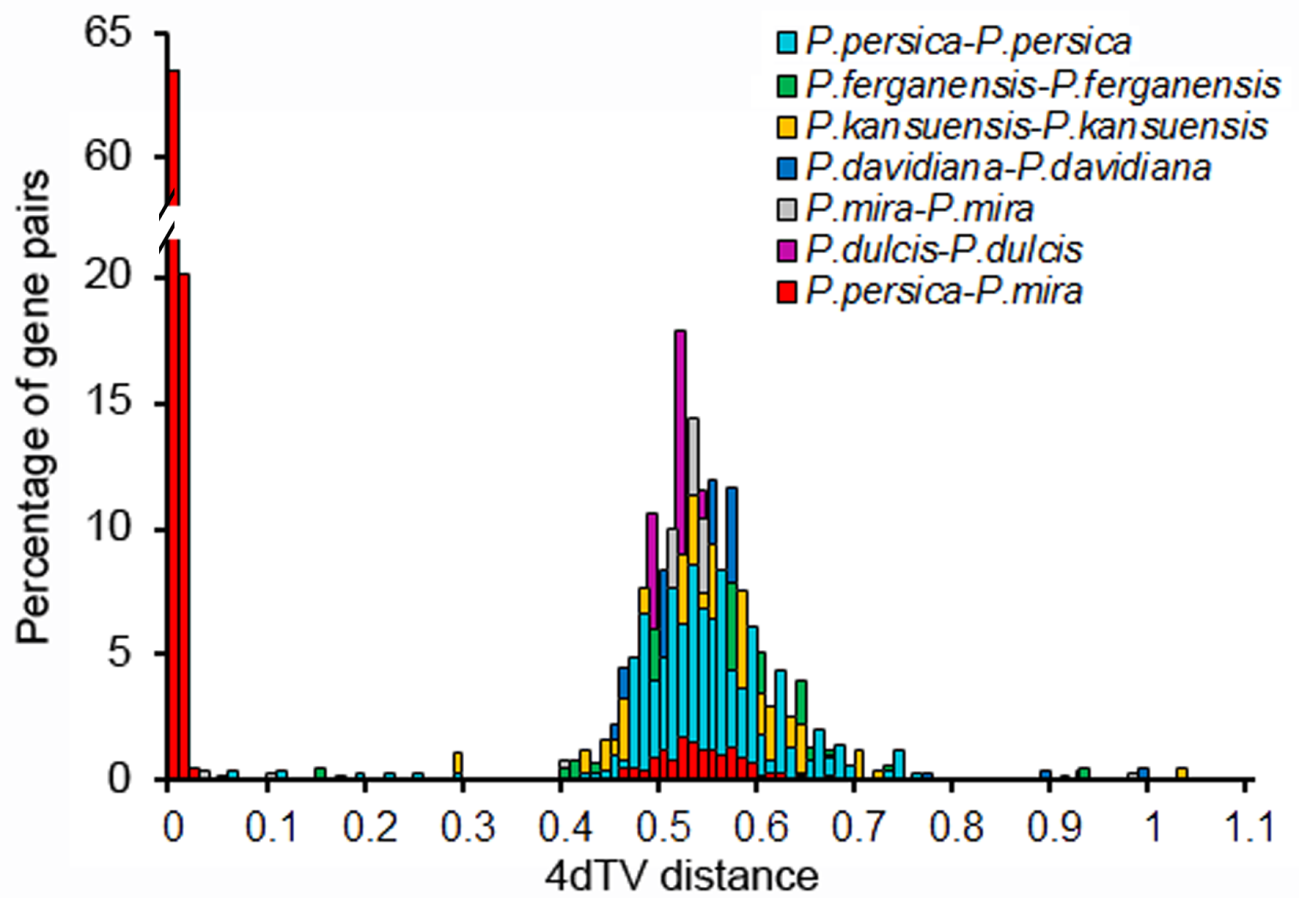

**Supplementary Figure 16 Percent of *P. davidiana*-specific contigs covered by reads from different *Prunus* species.**

The percent of reads in different varieties covered with contigs which assembled using unmapped sequence in P. davidiana aligned with P. mira

|  | 98% coverage | 95% coverage | 90% coverage | #REF! | #REF! |
| --- | --- | --- | --- | --- | --- |
| P. persica | 54.6028% | 58.0589% | 60.4717% | 100% | 100% |
| P. persica | 54.5267% | 57.7763% | 59.8196% |  |  |
| P. ferganensis | 50.1902% | 53.5485% | 55.907% |  |  |
| P. ferganensis | 42.3758% | 46.7775% | 50.2771% |  |  |
| P. kansuensis | 51.614% | 54.7223% | 57.059% |  |  |
| P. kansuensis | 44.3756% | 48.5925% | 52.1791% |  |  |
| P. dulcis | 63.2757% | 66.7754% | 69.0903% |  |  |
| P. tangutica | 57.6459% | 60.2761% | 62.3954% |  |  |
| P. armeniaca | 28.1926% | 33.4529% | 38.8219% |  |  |
| P. sibirica | 26.008% | 30.801% | 35.6374% |  |  |
| P. cerasifera | 26.1276% | 31.0292% | 35.9852% |  |  |
| P. salicina | 26.2689% | 31.051% | 35.9526% |  |  |
| P. avium | 17.9872% | 21.4977% | 25.9211% |  |  |
| P. mahaleb | 19.8565% | 23.8452% | 28.4317% |  |  |

**Supplementary Figure 17 Geographical distribution of *P. mira* (red circle), *P. davidiana* (yellow), and *P. dulcis* (orange) which originated in China.**

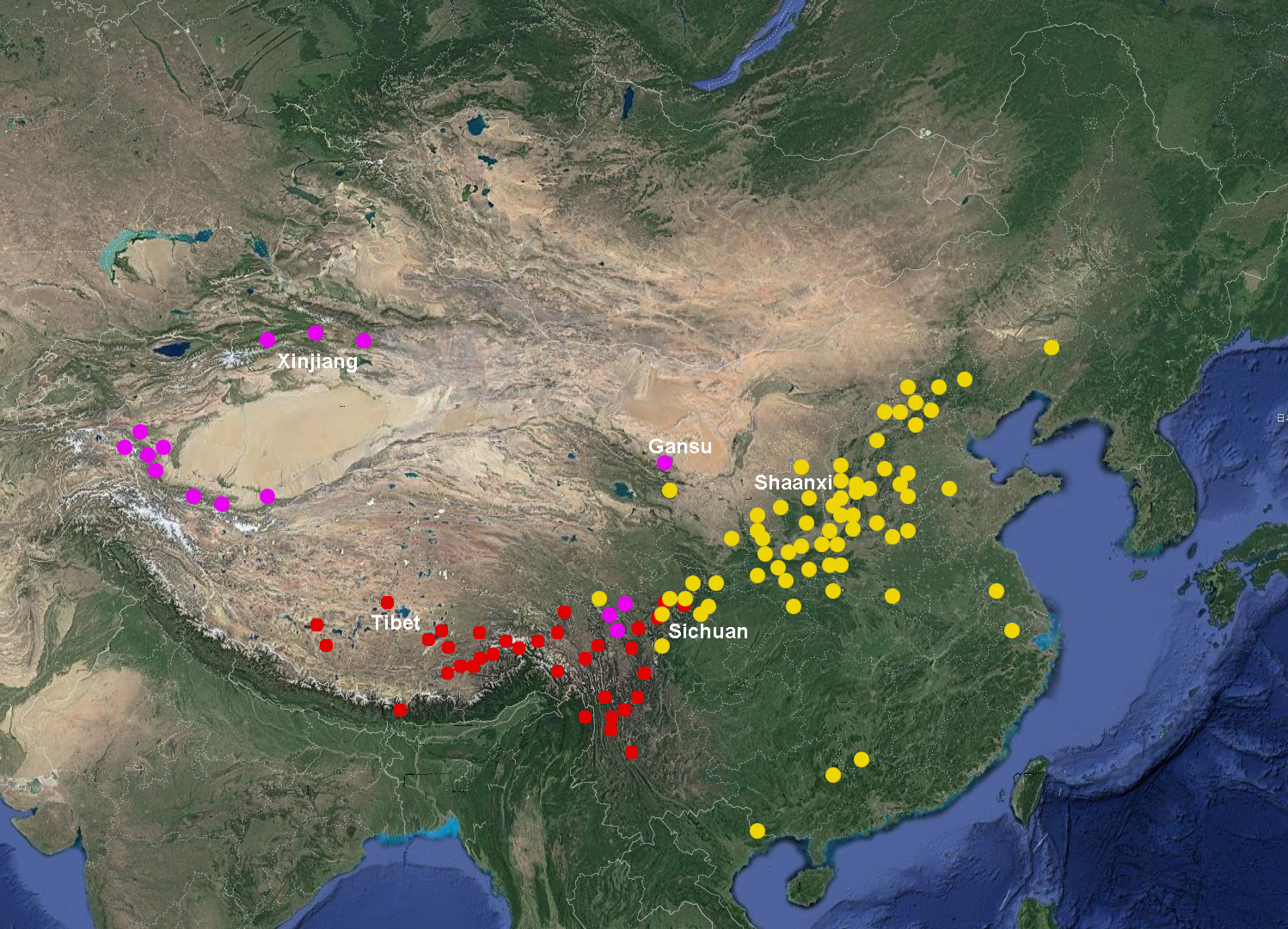

**Supplementary Figure 18 Distribution of resistance (*R*) genes across the 8 chromosomes in five peach species of peach and their overlaps with disease resistance QTLs.**

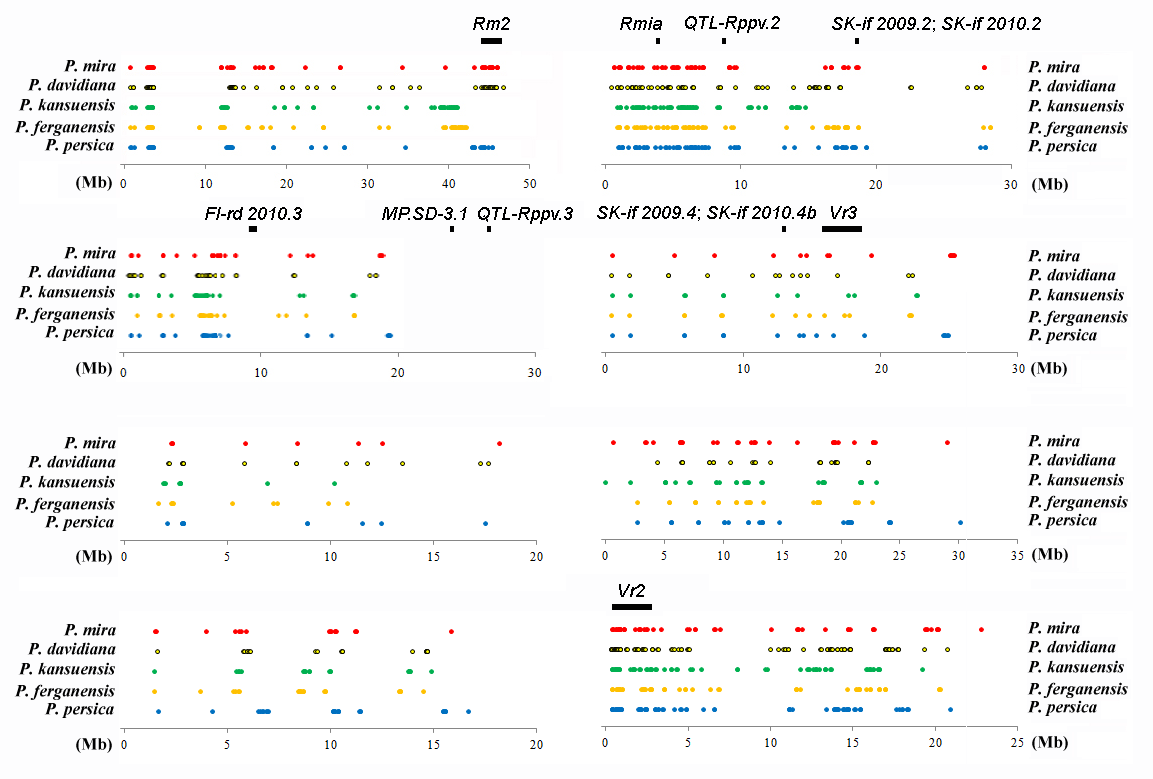

In the figure, ‘*Rm2*’ and ‘*MP.SD-3.1*’ indcated resistance gene or QTLs to green peach aphid which reported by Lambert et al. (2016) and Sauge et al. (2012), respectively, ‘*Rmia*’ indicated resistance to *Meloidogyne incognita* which reported by Duval et al. (2014), ‘*Rppv.2*’ and ‘*Rppv.3*’ indicated resistance QTLs to plum pox virus which reported by Cirilli et al. (2017), ‘*Vr2*’ and ‘*Vr3*’ indicated resistance to peach powdery mildew which reported by Pascal et al. (2017) and Donoso et al. (2016), ‘*SK-if 2009.2*’, ‘*SK-if 2010.2*’, ‘*Fl-rd 2010.3*’, ‘*SK-if 2009.4*’, and ‘*SK-if 2010.4b*’ indicated resistance QTLs to brown rot which reported by Pacheco et al. (2014). And the last number indicated the chromosome number which located that QTL.

**Supplementary Figure 19 KEGG pathways enriched in expanded and contracted gene families of the four wild peach species.**

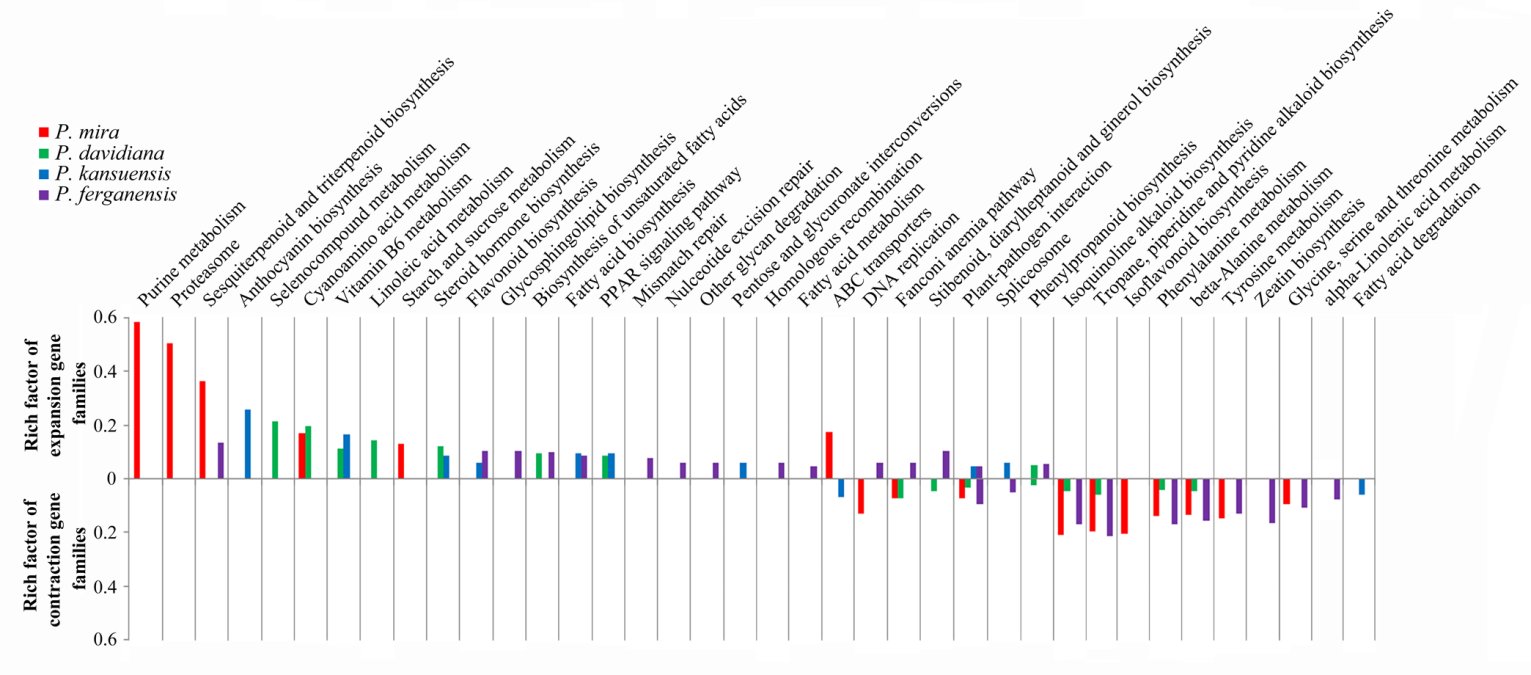

**Supplementary Figure 20 Phylogenetic tree of 32 accessions of *P. mira* (a) originating from regions with different altitudes (b) and the population structure (c) when K=2 and 3.**

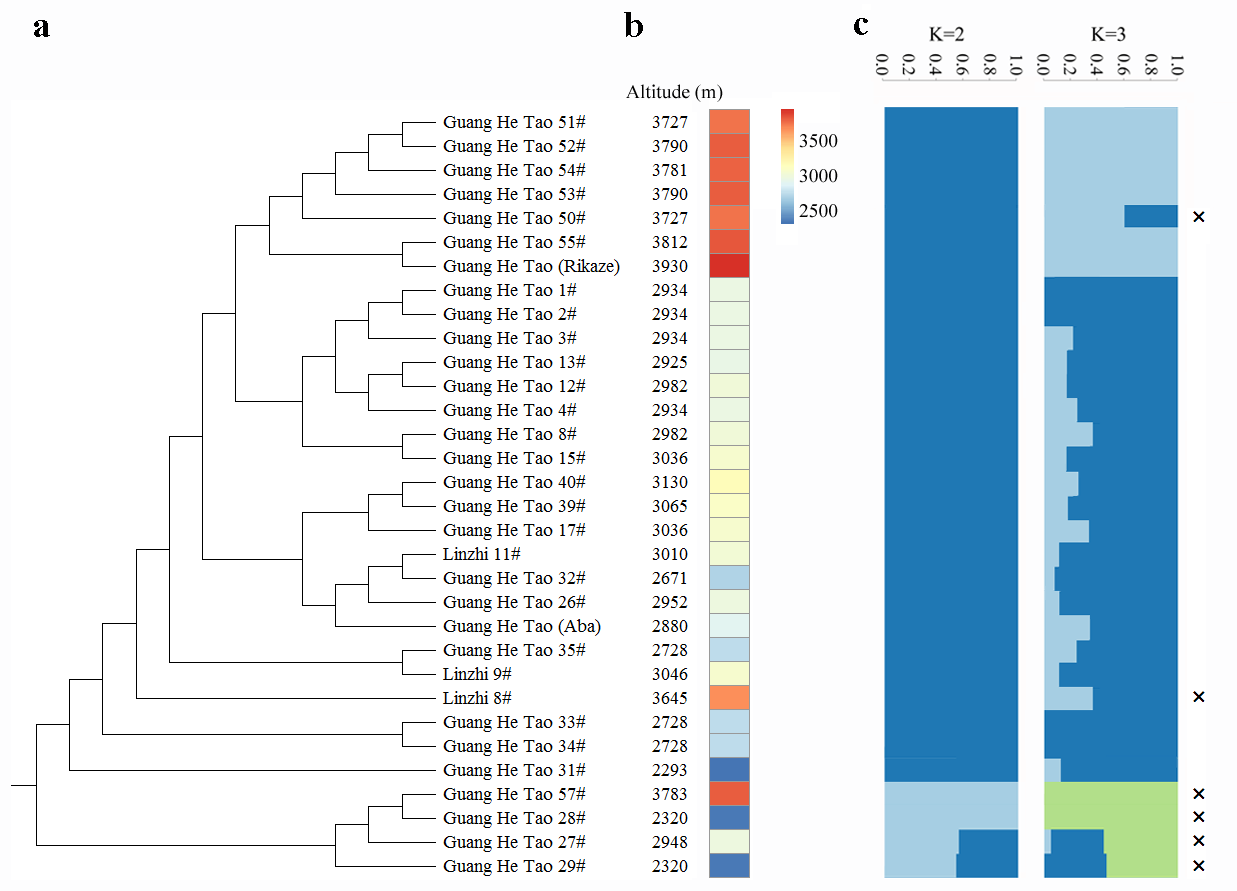

According to the phylogenetic tree and STRUCTURE analysis, three accessions thought to have not corresponded to its altitude categories, two reckoned as an admixture subgroup between high and low altitude subgroups, and one showed long distance with others were labelled with multiple sign and removed in the following analysis.

**Supplementary Figure 21 Two genome regions associated with high-altitude adaptation.**

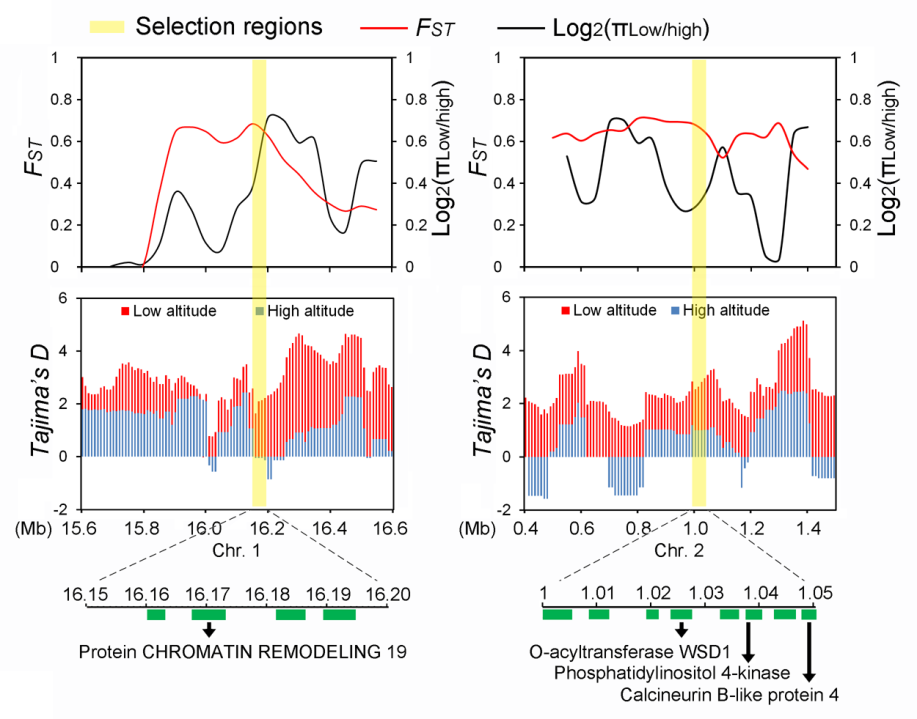

We also detected two genomic regions showing strong reduction of diversity and high differentiation. In the first region spanning from 16.15 Mb to 16.20 Mb of Chr. 1, *evm.model.Pm01.2222* encoded a protein Chromatin Remodeling 19 involved in DNA repair and heterochromatin organization. This finding is con­sistent with the observation that the high altitude adaptation of *C. himalaica* was related to DNA repair (Zhang et al., 2019). And in another region (Chr2: 1.00..1.05 Mb), *evm.model.Pm02.196* encoded O-acyltransferase WSD1-like protein which involved in drought response (Pan et al., 2018). *evm.model.Pm02.198* encoded Phosphatidylinositol 4-kinase gamma 3 responded to ABA and salt (McLoughlin et al., 2013). *evm.model.Pm02.200.2* encoded Calcineurin B-like protein 4 involved in salt response (Liu et al., 1997).
