## Supplementary table 6-24 for "Pan-genome analyses of peach and its wild relatives provide insights into the genetics of disease resistance and species adaptation"

**Supplementary Table 6 Validation of non-reference sequences using the Sanger technology.**

| Contig name | PCR product (bp) | The identity of PCR product to the reference genome v2.0 (%) | The identity of PCR product to the assembling sequences (%) |
| --- | --- | --- | --- |
| Contig3_A100 | 544 | 6.43 | 99.82 |
| Contig15_A10 | 317 | 49.21 | 100.00 |
| Contig17_A1 | 399 | 22.31 | 100.00 |
| Contig18_A11 | 571 | 5.95 | 100.00 |
| Contig35_A100 | 251 | 66.93 | 100.00 |
| Contig39_A66 | 345 | 78.55 | 97.97 |
| Contig40_A103 | 595 | - | 97.14 |
| Contig42_A104 | 557 | 29.62 | 100.00 |
| Contig55_A11 | 906 | 25.17 | 99.56 |
| Contig61_A11 | 373 | 58.71 | 100.00 |

**Supplementary Table 7 Genome survey of four wild peach species (kmer = 17).**

| ****Species**** | ****K-mer number**** | ****K-mer depth**** | ****Genome size (Mb)**** | ****Heterozygous ratio (%)**** | ****Repeat (%)**** |
| --- | --- | --- | --- | --- | --- |
| ***P. mira*** | 12,594,677,936 | 51 | 242.94 | 0.76 | 47.51 |
| ***P. davidiana*** | 22,358,636,352 | 93 | 237.29 | 1.10 | 45.09 |
| ***P. kansuensis*** | 14,480,733,904 | 60 | 238.06 | 0.56 | 46.07 |
| ***P. ferganensis*** | 13,266,273,280 | 55 | 237.24 | 0.53 | 45.09 |

Note: all values were calculated according to the curve of depth distribution of K-mer number.

**Supplementary Table 8 Summary of genome sequencing of four wild peach species.**

| Pair-end libraries | Insert size of libraries | *P. mira* | | *P. davidiana* | | *P. kansuensis* | | *P. ferganensis* | |
| --- | --- | --- | --- | --- | --- | --- | --- | --- | --- |
|  |  | Total data (Gb) | Coverage (×) | Total data (Gb) | Coverage (×) | Total data (Gb) | Coverage (×) | Total data (G) | Coverage (×) |
| Illumina | 230 bp | 14.99 | 61.70 | 14.30 | 60.26 | 13.8 | 52.08 | 12.9 | 54.38 |
|  | 500 bp | 15.43 | 63.51 | 15.63 | 65.87 | 5.6 | 21.13 | 5.2 | 21.92 |
|  | 2 Kb | 13.41 | 55.20 | 13.46 | 56.72 | 5.8 | 21.89 | 5.8 | 24.45 |
|  | 5 Kb | 6.57 | 27.04 | 9.71 | 40.92 | 6.8 | 25.66 | 6.2 | 26.13 |
|  | 10 Kb | 6.00 | 24.70 | 6.16 | 25.96 |  |  |  |  |
|  | 20 Kb | 2.57 | 10.58 | 3.31 | 13.95 |  |  |  |  |
| PacBio | 20 Kb | 13.93 | 57.34 |  |  |  |  |  |  |
| Hi-C | | 72.14 | 296.95 |  |  |  |  |  |  |
| Total | | 145.04 | 597.02 | 62.57 | 263.69 | 32.0 | 120.76 | 30.1 | 126.88 |

**Supplementary Table 9 Pseudochromosome lengths of the *P. mira* assembly.**

| Chr. | Length (bp) |
| --- | --- |
| Pm01 | 47,850,557 |
| Pm02 | 30,912,233 |
| Pm03 | 27,440,514 |
| Pm04 | 27,095,314 |
| Pm05 | 20,305,835 |
| Pm06 | 32,154,895 |
| Pm07 | 23,805,492 |
| Pm08 | 26,080,652 |

**Supplementary Table 10 BUSCO analysis of the genome assemblies of four wild peach species.**

| Species | BUSCO assessment result based on a total of 1440 searched BUSCO groups | | | |
| --- | --- | --- | --- | --- |
|  | **S (%)** | **D (%)** | **F (%)** | **M (%)** |
| *P. mira* | 90.3 | 5.2 | 1.1 | 3.4 |
| *P. davidiana* | 95.2 | 1.9 | 1.0 | 1.9 |
| *P. kansuensis* | 95.5 | 2.0 | 0.8 | 1.7 |
| *P. ferganensis* | 96.1 | 1.7 | 0.7 | 1.5 |

S: Complete and single-copy BUSCOs.

D: Complete and duplicated BUSCOs.

F: Fragmented BUSCOs.

M: Missing BUSCOs.

Note: More than 90% of embryophyta genes were detected in above table, indicating a good completeness of our assembly.

**Supplementary Table 11 Mapping statistics of RNA-Seq reads to the corresponding genome assemblies of four wild peach species.**

| Tissue | *P. mira* (%) | *P. davidiana* (%) | *P. kansuensis* (%) | *P. ferganensis* (%) |
| --- | --- | --- | --- | --- |
| **Fruit** | 81 | 84 | 84 | 82 |
| **Flower** | 82 | 81 | 86 | 86 |
| **Phloem** | 78 | 84 | 82 | 80 |
| **Leaf** | 80 | 85 | 79 | 84 |
| **Seed** | 83 | 83 | 82 | 78 |
| **Total** | 81 | 83 | 83 | 82 |

Note: the value indicates the percent of mapped reads to total sequenced reads in different accessions and tissues.

**Supplementary Table 12 Statistic of repeat sequences in the assemblies of four wild peach species.**

| **Category** | | ***P. mira*** | | ***P. davidiana*** | | ***P. kansuensis*** | | ***P. ferganensis*** | |
| --- | --- | --- | --- | --- | --- | --- | --- | --- | --- |
|  |  | **Length (bp)** | **% in genome** | **Length (bp)** | **% in genome** | **Length (bp)** | **% in genome** | **Length (bp)** | **% in genome** |
| **Total repeat** | | 119,715,215 | 47.43 | 89,224,920 | 40.46 | 79,198,752 | 38.41 | 78,640,823 | 38.44 |
| **Tandem repeat** | | 12,167,262 | 4.82 | 11,514,846 | 5.22 | 8,392,669 | 4.07 | 8,972,483 | 4.39 |
| **Interpersed repeat** | DNA | 37,914,818 | 15.02 | 24,130,082 | 10.94 | 20.979,380 | 10.18 | 20,578,568 | 10.06 |
|  | LINE | 3,891,578 | 1.54 | 3,748,431 | 1.7 | 3,180,866 | 1.54 | 2,603,654 | 1.27 |
|  | SINE | 249,458 | 0.1 | 108,512 | 0.05 | 33,696 | 0.02 | 199,374 | 0.1 |
|  | LTR | 64,192,379 | 25.43 | 52,666,789 | 23.88 | 49,624,236 | 24.07 | 48,104,838 | 23.51 |
|  | Unknown | 18,089,773 | 7.17 | 9,969,139 | 4.52 | 7,780,020 | 3.77 | 8,659,124 | 4.23 |

During the whole genome sequence, we identified 119.71 Mb repeat sequences using two approaches, De novo and homologous prediction, which represents 47.43% of the *P. mira* genome. The value showed higher than that of *P. persica* (37.14%, International Peach Genome Initiative, 2013), Kiwifruit (36%, Huang et al., 2013), and grape (41.4%, The French–Italian Public Consortium for Grapevine Genome Characterization, 2007) but lower than that of other fruit crops, such as apple (67.4%, Velasco et al., 2010), pear (53.1%, Wu et al., 2013), and soybean (53.9%, Xie et al., 2019). Moreover, we identified a total of 40.46%, 38.41%, and 38.44% of repeat sequence in *P. davidiana*, *P. kansuensis*, and *P. ferganensis*.

**Supplementary Table 13 Prediction of protein-coding genes in the genomes of four wild peach species.**

| Method used for gene annotation | | *P. mira* | *P. davidiana* | *P. kansuensis* | *P. ferganensis* |
| --- | --- | --- | --- | --- | --- |
| De novo | Augustus | 21,095 | 19,974 | 20,405 | 20,431 |
|  | Glimmer HMM | 34,844 | 32,855 | 32,781 | 31,713 |
|  | SNAP | 23,997 | 30,933 | 29,996 | 30,413 |
|  | Genscan | 19,424 | 18,945 | 19,144 | 19,473 |
|  | Geneid | 29,656 | 35,805 | 35,446 | 36,071 |
| Homolog | *P. persica* | 32,828 | 26,817 | 26,989 | 28,095 |
|  | *Pyrus bretschneideri* | 20,909 | 21,160 | 21,577 | 22,424 |
|  | *P. mume* | 31,304 | 22,089 | 22,637 | 23,670 |
|  | *Malus domestica* | 18,756 | 19,536 | 19,971 | 21,230 |
|  | *Fragaria vesca* | 19,715 | 21,930 | 22,420 | 23,109 |
|  | *Vitis vinifera* | 21,670 | 19,976 | 19,910 | 20,230 |
|  | *Arabidopsis thaliana* | 23,515 | 21,545 | 21,360 | 21,616 |
| RNA-seq | PASA | 66,856 | 62,870 | 57,243 | 61,714 |
|  | Cufflinks | 38,514 | 40,678 | 37,171 | 37,874 |
| Final set | | 28,943 | 26,527 | 26,297 | 27,431 |

Note: In the study, five software programs were used for gene structure *de novo* annotation, such as Augustus, Glimmer HMM, SNAP, Genscan, and Geneid. Meanwhile, *P. persica*, *Pyrus bretschneideri*, *P. mume*, *Malus domestica*, *Fragaria vesca*, *Vitis vinifera*, and *Arabidopsis thaliana* were used for gene annotation by homolog analysis. In addition, RNA-seq data of different tissues were also used for assistance annotation with different tools, such as PASA and Cufflinks. In the study, we identified 28,943, 26,527, 26,297, and 27,431 high-confidence protein-coding genes in *P. mira*, *P. davidiana*, *P. kansuensis*, and *P. ferganensis* genomes, respectively.

**Supplementary Table 14 Statistics of predicted protein-coding genes in four wild peach species compared to other species.**

| Species | Number | Average transcript length (bp) | Average CDS length (bp) | Average exons per gene | Average exon length (bp) | Average intron length (bp) |
| --- | --- | --- | --- | --- | --- | --- |
| *P. mira* | 28,943 | 2,653.19 | 1,100.00 | 4.70 | 233.95 | 419.58 |
| *P. davidiana* | 26,527 | 2,543.72 | 1,126.54 | 4.67 | 241.21 | 386.12 |
| *P. kansuensis* | 26,297 | 2,565.41 | 1,154.65 | 4.73 | 243.93 | 377.87 |
| *P. ferganensis* | 27,431 | 2,478.64 | 1,134.17 | 4.65 | 243.88 | 368.29 |
| *P. persica* | 27,796 | 2,457.59 | 1,205.92 | 4.85 | 248.58 | 325.00 |
| *P. mume* | 23,161 | 2,893.71 | 1,329.10 | 5.25 | 253.07 | 367.99 |
| *Pyrus bretschneideri* | 34,328 | 3,115.05 | 1,330.05 | 5.47 | 243.28 | 399.59 |
| *Malus domestica* | 45,442 | 3,528.27 | 1,205.82 | 4.82 | 250.4 | 608.66 |
| *Fragaria vesca* | 23,718 | 2,678.81 | 1,326.55 | 5.17 | 256.62 | 324.33 |
| *Vitis vinifera* | 29,825 | 4,737.52 | 1,097.24 | 4.76 | 230.73 | 968.99 |
| *Arabidopsis thaliana* | 27,379 | 1,871.96 | 1,214.84 | 5.13 | 236.93 | 159.16 |
| *Populus trichocarpa* | 40,045 | 2,345.98 | 1,121.48 | 4.74 | 236.83 | 327.68 |
| *Oryza sativa* | 34,608 | 2,202.50 | 998.81 | 3.83 | 260.77 | 425.06 |
| *Cucumis sativus* | 18,549 | 3,826.54 | 1,354.64 | 5.82 | 232.82 | 513.02 |

The gene transcripts had an average length of 2,457.59 ~ 2,653.19 bp, a mean coding sequence (CDS) size of 1,100.00 ~ 1205.92 bp and an average exon length of 233.95 ~ 248.58 bp in *P. persica* and its four wild related species. Among *Prunus*, the gene density was higher in *P. ferganensis* (13.41 genes/100 kb) than in *P. mira* (11.47 genes/100 kb), and shorter transcript but longer CDS were found in *P. persica* comparing with other *Prunus* indicating that the length of intron was decreased in peach and closest relatives.

**Supplementary Table 15 Statistics of gene functional annotation in the four wild peach species.**

| Database | | *P. mira* | | *P. davidiana* | | *P. kansuensis* | | *P. ferganensis* | |
| --- | --- | --- | --- | --- | --- | --- | --- | --- | --- |
|  |  | Number | Percent (%) | Number | Percent (%) | Number | Percent (%) | Number | Percent (%) |
| TrEMBL | | 27,155 | 93.8 | 25,006 | 94.3 | 24,762 | 94.2 | 25,546 | 93.1 |
| Swiss-Prot | | 21,016 | 72.6 | 19,522 | 73.9 | 19,523 | 74.2 | 19,987 | 72.9 |
| KEGG | | 20,459 | 70.7 | 18,589 | 70.1 | 18,649 | 70.9 | 19,132 | 69.7 |
| InterPro | All | 22,407 | 77.4 | 20,746 | 78.2 | 20,803 | 79.1 | 21,271 | 77.5 |
|  | Pfam | 20,498 | 70.8 | 19,485 | 73.5 | 19,596 | 74.5 | 20,006 | 72.9 |
|  | GO | 15,022 | 51.9 | 13,985 | 52.7 | 14,156 | 53.8 | 14,417 | 52.6 |
| Annotated | | 27,174 | 93.9 | 25,027 | 94.3 | 24,776 | 94.2 | 25,561 | 93.2 |
| Total | | 28,943 |  | 26,527 |  | 26,297 |  | 27,431 |  |

**Supplementary Table 16 Non-coding RNAs identified in genomes of four wild peach species.**

| Type | | *P. mira* | | *P. davidiana* | | *P. kansuensis* | | *P. ferganensis* | |
| --- | --- | --- | --- | --- | --- | --- | --- | --- | --- |
|  |  | Copy | Average length (bp) | Copy | Average length (bp) | Copy | Average length (bp) | Copy | Average length (bp) |
| miRNA | | 489 | 133.48 | 463 | 134.19 | 409 | 134.18 | 423 | 135.08 |
| tRNA | | 541 | 75.42 | 476 | 75.42 | 485 | 75.54 | 480 | 75.32 |
| rRNA |  | 195 | 322.58 | 90 | 261.41 | 49 | 189.33 | 51 | 186.65 |
|  | 18S | 42 | 984.21 | 36 | 456.61 | 15 | 367.53 | 20 | 290.25 |
|  | 28S | 92 | 146.65 | 15 | 148.67 | 12 | 111.67 | 8 | 131.5 |
|  | 5.8S | 26 | 156.85 | 11 | 151 | 4 | 139.25 | 4 | 157 |
|  | 5S | 35 | 114.17 | 28 | 114.21 | 18 | 103.72 | 19 | 107.05 |
| snRNA | snRNA | 449 | 115.66 | 374 | 115.46 | 358 | 115.4 | 340 | 117.16 |
|  | CD-box | 269 | 104.29 | 215 | 102.17 | 203 | 100.58 | 191 | 102.25 |
|  | HACA-box | 33 | 123.61 | 33 | 124 | 32 | 124 | 34 | 121.94 |
|  | Splicing | 146 | 134.71 | 125 | 135.94 | 122 | 137.7 | 114 | 140.6 |

The maximum number of microRNA, transfer RNA, ribosomal RNA, and small nuclear RNA were all annotated in *P. mira*.

**Supplementary Table 17 SNPs identified between genomes of each of the four wild species and *P. persica*.**

| Chromosome | *P. mira* | | *P. davidiana* | | *P. kansuensis* | | *P. ferganensis* | |
| --- | --- | --- | --- | --- | --- | --- | --- | --- |
|  | Number | Density  (bp/SNP) | Number | Density  (bp/SNP) | Number | Density  (bp/SNP) | Number | Density  (bp/SNP) |
| Chr.1 | 1074908 | 44.52 | 953537 | 50.18 | 724620 | 66.04 | 147499 | 324.42 |
| Chr.2 | 589134 | 51.61 | 631709 | 48.13 | 536954 | 56.63 | 193810 | 156.88 |
| Chr.3 | 556977 | 49.14 | 598030 | 45.76 | 498554 | 54.89 | 97353 | 281.12 |
| Chr.4 | 514925 | 50.19 | 501170 | 51.57 | 413535 | 62.49 | 151340 | 170.76 |
| Chr.5 | 353319 | 52.35 | 366352 | 50.49 | 284926 | 64.92 | 53847 | 343.50 |
| Chr.6 | 609415 | 50.49 | 630443 | 48.80 | 474182 | 64.88 | 203544 | 151.16 |
| Chr.7 | 460516 | 48.62 | 466398 | 48.00 | 362728 | 61.72 | 80535 | 278.00 |
| Chr.8 | 482089 | 46.83 | 514470 | 43.88 | 409979 | 55.06 | 108802 | 207.48 |
| Assembly | 4,348,360 |  | 4,440,297 |  | 3,491,940 |  | 992,783 |  |
| Reads | 3,589,992 |  | 3,828,237 |  | 2,849,820 |  | 669,084 |  |
| Combined | 4,683,941 |  | 4,696,627 |  | 3,744,223 |  | 1,062,698 |  |

Note: ‘Assembly’ indicated the SNPs detected based on genome *de novo* assembling sequence. ‘Reads’ indicated the SNPs detected using clean reads. ‘Combined’ indicated the SNPs which calculated using the above two methods.

**Supplementary Table 18 Small indels (<50 bp) identified between genomes of each of the four wild species and *P. persica*.**

| Species | Insertion | | | Deletion | | | Total gene |
| --- | --- | --- | --- | --- | --- | --- | --- |
|  | Length (bp) | Number | Affected gene number | Length (bp) | Number | Affected gene number |  |
| *P. mira* | 1,181,090 | 266,143 | 3,339 | 1,098,209 | 252,723 | 2,272 | 5,438 |
| *P. davidiana* | 1,169,377 | 260,257 | 3,053 | 976,464 | 241,439 | 1,979 | 4,807 |
| *P. kansuensis* | 939,524 | 217,306 | 2,555 | 973,999 | 218,866 | 1,998 | 4,352 |
| *P. ferganensis* | 249,906 | 56,726 | 770 | 261,277 | 58,802 | 488 | 1,217 |

**Supplementary Table 19 Genome structural variants (≥ 50 bp) between the four wild species and *P. persica*.**

| Species | Insertion | | | Inversion | | | Deletion | | | Total gene |
| --- | --- | --- | --- | --- | --- | --- | --- | --- | --- | --- |
|  | Length (bp) | Number | Affected gene number | Length (bp) | Number | Affected gene number | Length (bp) | Number | Affected gene number |  |
| *P. mira* | 245,037 | 3,526 | 122 | 1,749,938 | 2,083 | 229 | 191,718 | 2,809 | 62 | 410 |
| *P. davidiana* | 248,398 | 3,594 | 129 | 1,427,987 | 1,833 | 142 | 164,055 | 2,396 | 47 | 315 |
| *P. kansuensis* | 200,138 | 2,896 | 88 | 1,923,118 | 1,728 | 282 | 191,500 | 2,810 | 69 | 436 |
| *P. ferganensis* | 82,013 | 1,151 | 34 | 665,729 | 446 | 79 | 62,120 | 878 | 24 | 137 |

**Supplementary Table 20 Statistics of copy number variations between the four wild species and *P. persica.***

| Species | Deletion | | | Duplication | | | Total gene |
| --- | --- | --- | --- | --- | --- | --- | --- |
|  | Length (bp) | Number | Affected gene number | Length (bp) | Number | Affected gene number |  |
| *P. mira* | 46,535,100 | 5,426 | 2,353 | 10,350,600 | 1,461 | 1,307 | 3,610 |
| *P. davidiana* | 44,448,600 | 6,145 | 2,052 | 5,036,000 | 945 | 677 | 2,703 |
| *P. kansuensis* | 38,339,300 | 4,534 | 1,857 | 9,016,900 | 1,260 | 1,034 | 2,873 |
| *P. ferganensis* | 16,518,400 | 2,757 | 842 | 10,136,700 | 1,396 | 434 | 1,270 |

**Supplementary Table 21 Variations in the promoter and mRNA regions of *R* genes on Chr. 2 (5-7 Mb) that were specific to *P. kansuensis*.**

| *R* gene | | | Variations | | | | | |
| --- | --- | --- | --- | --- | --- | --- | --- | --- |
| Accession name | Start position (bp) | End position (bp) | Start position (bp) | End position (bp) | Genotype | Genotype | Variation type | Annotation |
| *Prupe.2G045200* | 5054083 | 5060312 | 5055163 | 5055163 | a | - | DEL | Exon |
|  |  |  | 5058166 | 5058166 | - | t | INS | Intron |
| *Prupe.2G046000* | 5254014 | 5257565 | 5254422 | 5254422 | t | - | DEL | Exon |
|  |  |  | 5255761 | 5255761 | - | caa | INS | Exon |
|  |  |  | 5255768 | 5255768 | - | tc | INS | Exon |
|  |  |  | 5255774 | 5255774 | - | t | INS | Exon |
|  |  |  | 5255780 | 5255780 | - | gca | INS | Exon |
| *Prupe.2G046300* | 5308864 | 5313947 | 5308309 | 5308309 | - | act | INS | Promoter |
| *Prupe.2G046400* | 5370808 | 5377086 | 5371512 | 5371512 | - | g | INS | 5'UTR |
|  |  |  | 5376152 | 5376152 | c | - | DEL | Intron |
|  |  |  | 5376162 | 5376162 | - | taatg | INS | Intron |
| *Prupe.2G046700* | 5408587 | 5413667 | 5408165 | 5408165 | a | - | DEL | Promoter |
|  |  |  | 5408287 | 5408287 | - | ttggtgacgtt | INS | Promoter |
|  |  |  | 5413439 | 5413439 | - | t | INS | 3'UTR |
|  |  |  | 5413441 | 5413441 | - | attg | INS | 3'UTR |
|  |  |  | 5413442 | 5413442 | - | a | INS | 3'UTR |
| *Prupe.2G046900* | 5445798 | 5452769 | 5444796 | 5444796 | a | - | DEL | Promoter |
|  |  |  | 5445199 | 5445199 | - | c | INS | Promoter |
|  |  |  | 5445363 | 5445363 | - | ttt | INS | Promoter |
|  |  |  | 5446932 | 5446932 | - | g | INS | 5'UTR |
|  |  |  | 5451241 | 5451241 | - | ctctct | INS | Intron |
|  |  |  | 5451461 | 5451464 | catg | - | DEL | Intron |
|  |  |  | 5451702 | 5451707 | tatgat | - | DEL | Intron |
| *Prupe.2G051100* | 5902195 | 5905326 |  |  |  |  |  |  |
| *Prupe.2G052200* | 6060321 | 6064679 | 6064779 | 6064779 | - | at | INS | Promoter |
|  |  |  | 6065340 | 6065340 | - | tataaac | INS | Promoter |
| *Prupe.2G053600* | 6212618 | 6217340 | 6212387 | 6212422 | tcttgcgtccaactttgccgaacccatccccatgcg | - | DEL | Promoter |
|  |  |  | 6212424 | 6212425 | gg | - | DEL | Promoter |
|  |  |  | 6217200 | 6217201 | ca | - | DEL | 3'UTR |
| *Prupe.2G053700* | 6258789 | 6263801 | 6260264 | 6260264 | - | c | INS | Exon |
|  |  |  | 6260269 | 6260269 | c | - | DEL | Exon |
| *Prupe.2G053800* | 6279090 | 6292484 | 6285258 | 6285258 | a | - | DEL | Intron |
|  |  |  | 6285437 | 6285438 | aa | - | DEL | Intron |
|  |  |  | 6285531 | 6285532 | aa | - | DEL | Intron |
|  |  |  | 6285583 | 6285583 | - | tata | INS | Intron |
|  |  |  | 6285666 | 6285703 | tctctctctctctctctctctctctctctctctctctc | - | DEL | Intron |
|  |  |  | 6287398 | 6287402 | aaaaa | - | DEL | 3'UTR |
|  |  |  | 6287620 | 6287621 | aa | - | DEL | 3'UTR |
|  |  |  | 6287910 | 6287910 | - | taagaaaggtggaaccctgagt | INS | 3'UTR |
|  |  |  | 6287911 | 6287911 | - | gaag | INS | 3'UTR |
|  |  |  | 6288114 | 6288115 | aa | - | DEL | 3'UTR |
|  |  |  | 6290154 | 6290154 | a | - | DEL | Intron |
| *Prupe.2G053900* | 6309628 | 6311069 | 6308634 | 6308634 | - | c | INS | Promoter |
|  |  |  | 6308713 | 6308713 | - | aa | INS | Promoter |
|  |  |  | 6309001 | 6309001 | - | tctc | INS | Promoter |
|  |  |  | 6309006 | 6309006 | - | a | INS | Promoter |
|  |  |  | 6309019 | 6309019 | - | c | INS | Promoter |
| *Prupe.2G054000* | 6311430 | 6313309 |  |  |  |  |  |  |
| *Prupe.2G054100* | 6330046 | 6330862 |  |  |  |  |  |  |
| *Prupe.2G054300* | 6369296 | 6375443 | 6368554 | 6368554 | - | c | INS | Promoter |
|  |  |  | 6369021 | 6369021 | - | ttta | INS | Promoter |
|  |  |  | 6372754 | 6372754 | t | - | DEL | Exon |
|  |  |  | 6375260 | 6375260 | - | tt | INS | 3'UTR |
| *Prupe.2G054500* | 6428085 | 6431952 | 6429759 | 6429759 | - | a | INS | Promoter |
| *Prupe.2G054700* | 6460427 | 6466614 |  |  |  |  |  |  |
| *Prupe.2G055200* | 6542068 | 6547629 | 6541428 | 6541429 | gt | - | DEL | Promoter |
|  |  |  | 6541858 | 6541858 | a | - | DEL | Promoter |
|  |  |  | 6543595 | 6543596 | aa | - | DEL | Intron |
| *Prupe.2G055500* | 6600515 | 6601705 |  |  |  |  |  |  |
| *Prupe.2G055600* | 6636553 | 6641016 | 6637199 | 6637199 | - | t | INS | Intron |
| *Prupe.2G055700* | 6647990 | 6652171 | 6646588 | 6646588 | t | - | DEL | Promoter |
|  |  |  | 6649655 | 6649655 | - | cga | INS | Exon |
|  |  |  | 6649677 | 6649679 | aat | - | DEL | Exon |
|  |  |  | 6650005 | 6650005 | a | - | DEL | Exon |
|  |  |  | 6650016 | 6650016 | - | g | INS | Exon |
|  |  |  | 6650053 | 6650053 | - | g | INS | Exon |
|  |  |  | 6650059 | 6650059 | - | a | INS | Exon |
|  |  |  | 6650060 | 6650060 | - | a | INS | Exon |
|  |  |  | 6651488 | 6651488 | - | tg | INS | Intron |
|  |  |  | 6651512 | 6651519 | gttatatg | - | DEL | Intron |
|  |  |  | 6652031 | 6652031 | g | - | DEL | Exon |
| *Prupe.2G056100* | 6680147 | 6684535 | 6678550 | 6678550 | - | gcc | INS | Promoter |
|  |  |  | 6678837 | 6678848 | gccacacgcgcc | - | DEL | Promoter |
|  |  |  | 6679306 | 6679306 | - | c | INS | Promoter |
|  |  |  | 6681306 | 6681310 | ttggt | - | DEL | Exon |
|  |  |  | 6684496 | 6684500 | atggt | - | DEL | 3'UTR |
| *Prupe.2G057000* | 6859284 | 6859837 | 6860515 | 6860515 | - | a | INS | Promoter |
| *Prupe.2G057100* | 6901141 | 6907913 | 6902540 | 6902540 | a | - | DEL | Exon |
|  |  |  | 6902542 | 6902542 | - | c | INS | Exon |
|  |  |  | 6905079 | 6905079 | - | aca | INS | Exon |
|  |  |  | 6905516 | 6905516 | - | attttagca | INS | Exon |
|  |  |  | 6905747 | 6905747 | - | cctccatgtgt | INS | Intron |
|  |  |  | 6906043 | 6906043 | - | agacacattcaatgaatc | INS | Intron |

**Supplementary Table 22 Statistics of resistance genes in the four wild peach species.**

| Categories | | *P. mira* | *P. davidiana* | *P. kasuensis* | *P. ferganensis* |
| --- | --- | --- | --- | --- | --- |
| CNL | CC-NBS | 4 | 9 | 6 | 9 |
|  | CC-NBS-LRR | 39 | 56 | 44 | 57 |
| TNL | TIR-NBS | 20 | 21 | 16 | 22 |
|  | TIR-NBS-LRR | 54 | 86 | 88 | 79 |
| Unclassified | NBS | 76 | 52 | 44 | 40 |
|  | NBS-LRR | 115 | 110 | 124 | 108 |
|  | NBS-LRR-TIR | 0 | 0 | 0 | 0 |
|  | NBS-LRR-NBS-LRR | 0 | 0 | 0 | 1 |
|  | TIR-NBS-LRR-TIR | 0 | 0 | 0 | 0 |
|  | TIR-CC-NBS-LRR | 1 | 1 | 0 | 0 |
|  | TIR-NBS-LRR-NBS-LRR | 0 | 2 | 0 | 3 |
|  | TIR-NBS-LRR-TIR-NBS-LRR | 1 | 1 | 0 | 1 |
|  | TIR-NBS-TIR-NBS-LRR | 0 | 1 | 1 | 0 |
|  | LRR-TIR-NBS-LRR | 0 | 0 | 0 | 0 |
| Total genes | | 310 | 339 | 323 | 320 |
| Total types | | 8 | 10 | 7 | 9 |

**Supplementary Table 23 Genes selected between the two subgroups of *P. mira* which originated from high- and low-altitude regions.**

| Gene ID | Function annotated by Swiss-Prot database |
| --- | --- |
| *evm.model.Pm07.1801* | NA |
| *evm.model.Pm07.1802.1* | Protein PHR1-LIKE 1 |
| *evm.model.Pm07.1803* | UBP1-associated protein 2C |
| *evm.model.Pm07.1804* | NA |
| *evm.model.Pm07.1805* | NA |
| *evm.model.Pm07.1806* | NA |
| *evm.model.Pm07.1807* | NA |
| *evm.model.Pm07.1827* | Probable inactive leucine-rich repeat receptor-like protein kinase At1g66830 |
| *evm.model.Pm07.1828* | WD repeat-containing protein 55 |
| *evm.model.Pm07.1829* | OTU domain-containing protein At3g57810 |
| *evm.model.Pm07.1830.2* | Protein cornichon homolog 1 |
| *evm.model.Pm07.1831* | Transcription factor bHLH48 |
| *evm.model.Pm07.1832* | Exo-poly-alpha-D-galacturonosidase |
| *evm.model.Pm07.588* | 1-aminocyclopropane-1-carboxylate synthase 7 |
| *evm.model.Pm07.589* | Xaa-Pro dipeptidase |
| *evm.model.Pm07.590* | Probable protein arginine N-methyltransferase 1 |
| *evm.model.Pm07.591* | NA |
| *evm.model.Pm07.592* | NA |
| *evm.model.Pm07.593* | NA |
| *evm.model.Pm07.594* | Pentatricopeptide repeat-containing protein At2g27610 |
| *evm.model.Pm07.595* | Pentatricopeptide repeat-containing protein At4g13650 |
| *evm.model.Pm07.596.1* | Pollen-specific protein SF21 |
| *evm.model.Pm07.597* | Inorganic pyrophosphatase 3 |
| *evm.model.Pm07.598* | Graves disease carrier protein |
| *evm.model.Pm07.599* | Pentatricopeptide repeat-containing protein |
| *evm.model.Pm07.600* | Serine acetyltransferase 5 |
| *evm.model.Pm07.601* | NA |
| *evm.model.Pm07.602* | Inactive protein kinase SELMODRAFT_444075 |
| *evm.model.Pm07.603* | Probable acyl-[acyl-carrier-protein]--UDP-N-acetylglucosamine O-acyltransferase |
| *evm.model.Pm07.604* | Transcription factor DIVARICATA |
| *evm.model.Pm07.605* | K(+) efflux antiporter 4 |
| *evm.model.Pm07.606* | 1-aminocyclopropane-1-carboxylate oxidase 1 |
| *evm.model.Pm07.607* | Fanconi anemia group E protein |
| *evm.model.Pm07.608* | Tetraspanin-2 |
| *evm.model.Pm07.609* | Malonyl-[acyl-carrier protein] O-methyltransferase |
| *evm.model.Pm07.669* | Uncharacterized membrane protein C776.05 |
| *evm.model.Pm07.670* | NA |
| *evm.model.Pm07.671* | Ankyrin repeat-containing protein At2g01680 |
| *evm.model.Pm07.672* | Ankyrin repeat-containing protein At2g01680 |
| *evm.model.Pm07.673* | PHD finger protein ALFIN-LIKE 5 |
| *evm.model.Pm07.674.2* | Protein bem46 |
| *evm.model.Pm07.675* | Nuclear transcription factor Y subunit A-3 |
| *evm.model.Pm07.676* | Phospholipid:diacylglycerol acyltransferase 1 |
| *evm.model.Pm07.677* | Phospholipid:diacylglycerol acyltransferase 1 |
| *evm.model.Pm07.678* | Protein Brevis radix-like 4 |
| *evm.model.Pm09.1* | Calmodulin |
| *evm.model.Pm09.2* | Vacuolar-processing enzyme |
| *evm.model.Pm09.4* | NA |
| *evm.model.Pm09.5* | Vacuolar-processing enzyme |
| *evm.model.Pm05.1684* | Probable myosin-binding protein 6 |
| *evm.model.Pm05.1685* | Serine/threonine-protein phosphatase PP2A-5 catalytic subunit |
| *evm.model.Pm05.1686* | Germin-like protein subfamily T member 2 |
| *evm.model.Pm05.1687* | 5-formyltetrahydrofolate cyclo-ligase, mitochondrial |
| *evm.model.Pm05.1688* | Germin-like protein subfamily T member 2 |
| *evm.model.Pm05.1689* | Germin-like protein subfamily T member 2 |
| *evm.model.Pm05.1690* | Protein TPLATE |
| *evm.model.Pm05.1691* | Germin-like protein subfamily T member 2 |
| *evm.model.Pm05.1692* | Transcription factor bHLH79 |
| *evm.model.Pm05.1693* | Dynein light chain, cytoplasmic |
| *evm.model.Pm05.1694* | NA |
| *evm.model.Pm05.1695* | Transcription factor MYB35 |
| *evm.model.Pm05.1696* | Probable beta-1,3-galactosyltransferase 19 |
| *evm.model.Pm05.1732* | Nuclear ribonuclease Z |
| *evm.model.Pm05.1733* | NA |
| *evm.model.Pm05.1734* | Delta-1-pyrroline-5-carboxylate dehydrogenase 12A1 |
| *evm.model.Pm05.1735* | Probable WRKY transcription factor 72 |
| *evm.model.Pm05.1736* | Ubiquitin-conjugating enzyme |
| *evm.model.Pm05.1737* | F-box protein SKIP23 |
| *evm.model.Pm05.1738* | NA |
| *evm.model.Pm05.1740* | Protein IQ-DOMAIN 31 |
| *evm.model.Pm05.1741* | ABC transporter A family member 2 |
| *evm.model.Pm05.257* | Disease resistance protein RPM1 |
| *evm.model.Pm05.258* | NA |
| *evm.model.Pm05.260* | NA |
| *evm.model.Pm05.272* | Eukaryotic translation initiation factor 3 subunit J |
| *evm.model.Pm05.273* | Alpha/beta hydrolase domain-containing protein 17C |
| *evm.model.Pm06.1721* | NA |
| *evm.model.Pm06.1722* | Nuclear transcription factor Y subunit B-3 |
| *evm.model.Pm06.1723* | Protein CPR-5 |
| *evm.model.Pm06.1724.1* | NA |
| *evm.model.Pm01.2221* | Transcription factor bHLH111 |
| *evm.model.Pm01.2222* | Protein CHROMATIN REMODELING 19 |
| *evm.model.Pm01.2223* | Protein indeterminate-domain 4, chloroplastic |
| *evm.model.Pm01.2224* | Protein indeterminate-domain 5, chloroplastic |
| *evm.model.Pm01.2334* | Putative clathrin assembly protein At2g01600 |
| *evm.model.Pm01.2335* | Probable NAD(P)H dehydrogenase subunit CRR3, chloroplastic |
| *evm.model.Pm01.2336* | U-box domain-containing protein 12 |
| *evm.model.Pm01.2337.1* | NAD(P)H-quinone oxidoreductase subunit L, chloroplastic |
| *evm.model.Pm01.2338* | Transmembrane protein 214-B |
| *evm.model.Pm01.2340* | Protein BCCIP homolog |
| *evm.model.Pm01.2341* | Protein BCCIP homolog |
| *evm.model.Pm01.2342* | DELLA protein GAIP-B |
| *evm.model.Pm01.2343* | Uncharacterized protein At4g22758 |
| *evm.model.Pm01.2360* | NA |
| *evm.model.Pm01.2361* | Mitogen-activated protein kinase 12 |
| *evm.model.Pm01.2362* | NA |
| *evm.model.Pm01.2363* | Serine carboxypeptidase-like 50 |
| *evm.model.Pm01.2364* | Serine carboxypeptidase-like 50 |
| *evm.model.Pm01.2365* | ATP-dependent DNA helicase RecG |
| *evm.model.Pm01.2366* | Homeobox-leucine zipper protein HOX3 |
| *evm.model.Pm01.2367* | Auxin efflux carrier component 3 |
| *evm.model.Pm01.2368* | Probable aquaporin NIP-type |
| *evm.model.Pm01.2369* | Protein WVD2-like 1 |
| *evm.model.Pm01.2371* | NA |
| *evm.model.Pm01.2372* | NA |
| *evm.model.Pm01.2373* | DnaJ homolog subfamily B member 8 |
| *evm.model.Pm01.2374* | NA |
| *evm.model.Pm01.2375* | NA |
| *evm.model.Pm01.2395* | ELL-associated factor 1 |
| *evm.model.Pm01.2396* | ABC transporter G family member 7 |
| *evm.model.Pm01.2397* | NA |
| *evm.model.Pm01.2398* | NA |
| *evm.model.Pm01.2399* | Uncharacterized transporter C5D6.04 |
| *evm.model.Pm01.2400* | Probable ribose-5-phosphate isomerase 2 |
| *evm.model.Pm01.2401* | NA |
| *evm.model.Pm01.2402* | Putative fasciclin-like arabinogalactan protein 20 |
| *evm.model.Pm01.2403* | Probable E3 ubiquitin ligase SUD1 |
| *evm.model.Pm01.2404* | Sulfhydryl oxidase 2 |
| *evm.model.Pm01.2405* | Repressor of RNA polymerase III transcription MAF1 homolog |
| *evm.model.Pm01.2406* | NA |
| *evm.model.Pm01.3424* | Rhomboid-like protein 14, mitochondrial |
| *evm.model.Pm01.3425* | F-box/kelch-repeat protein At1g16250 |
| *evm.model.Pm01.3426* | Putative F-box protein At3g25750 |
| *evm.model.Pm01.3428* | GPI ethanolamine phosphate transferase 3 |
| *evm.model.Pm01.3429* | Protein NRT1/ PTR FAMILY 4.3 |
| *evm.model.Pm01.3431* | Pectin acetylesterase 10 |
| *evm.model.Pm01.3432* | Transcription factor HY5-like |
| *evm.model.Pm01.3433.1* | NA |
| *evm.model.Pm01.3434* | NA |
| *evm.model.Pm01.3435.1* | Chromatin structure-remodeling complex protein BSH |
| *evm.model.Pm01.3436* | NA |
| *evm.model.Pm01.3437* | NA |
| *evm.model.Pm01.3517* | Peroxisomal membrane protein 13 |
| *evm.model.Pm01.3518* | ABC transporter F family member 4 |
| *evm.model.Pm01.3520* | Protein NLP8 |
| *evm.model.Pm01.3522.1* | TBC1 domain family member 15 |
| *evm.model.Pm01.3523* | Probable peroxidase 26 |
| *evm.model.Pm01.3610* | Protein EFR3 homolog cmp44E |
| *evm.model.Pm01.3611* | ATPase family AAA domain-containing protein At1g05910 |
| *evm.model.Pm01.3612* | NA |
| *evm.model.Pm01.3631* | Plant UBX domain-containing protein 11 |
| *evm.model.Pm01.3632* | NA |
| *evm.model.Pm01.3633* | Plant UBX domain-containing protein 11 |
| *evm.model.Pm01.3634* | Biotin synthase |
| *evm.model.Pm01.3636.1* | PTI1-like tyrosine-protein kinase 3 |
| *evm.model.Pm01.3639* | Gem-associated protein 2 |
| *evm.model.Pm01.3640* | Putative disease resistance protein RGA3 |
| *evm.model.Pm01.3743* | Vacuolar amino acid transporter 1 |
| *evm.model.Pm01.3744* | Eukaryotic translation initiation factor 3 subunit H |
| *evm.model.Pm01.3745* | Probably inactive leucine-rich repeat receptor-like protein kinase At3g28040 |
| *evm.model.Pm01.3746* | Uncharacterized WD repeat-containing protein C2A9.03 |
| *evm.model.Pm01.3747* | NA |
| *evm.model.Pm01.3748* | Protein ROOT PRIMORDIUM DEFECTIVE 1 |
| *evm.model.Pm03.2078* | NA |
| *evm.model.Pm03.2079* | Small subunit processome component 20 homolog |
| *evm.model.Pm03.2080* | Heat shock 70 kDa protein, mitochondrial |
| *evm.model.Pm03.2082* | NA |
| *evm.model.Pm03.2083* | Pleiotropic drug resistance protein 1 |
| *evm.model.Pm02.10* | Growth-regulating factor 5 |
| *evm.model.Pm02.100* | NAC domain-containing protein 62 |
| *evm.model.Pm02.101* | IAA-amino acid hydrolase ILR1-like 5 |
| *evm.model.Pm02.103* | Heavy metal-associated isoprenylated plant protein 26 |
| *evm.model.Pm02.104* | NA |
| *evm.model.Pm02.105* | Peroxidase 5 |
| *evm.model.Pm02.106* | Nascent polypeptide-associated complex subunit alpha-like protein 2 |
| *evm.model.Pm02.107.2* | NA |
| *evm.model.Pm02.108* | 1-aminocyclopropane-1-carboxylate oxidase 1 |
| *evm.model.Pm02.11* | GPCR-type G protein 1 |
| *evm.model.Pm02.12* | Bromodomain adjacent to zinc finger domain protein 2B |
| *evm.model.Pm02.13* | NA |
| *evm.model.Pm02.14* | Aliphatic (R)-hydroxynitrile lyase |
| *evm.model.Pm02.193* | Histone-lysine N-methyltransferase ATX5 |
| *evm.model.Pm02.194* | Zinc finger protein CONSTANS-LIKE 12 |
| *evm.model.Pm02.195* | Protein BOBBER 1 |
| *evm.model.Pm02.196* | O-acyltransferase WSD1 |
| *evm.model.Pm02.197* | Probable pectinesterase/pectinesterase inhibitor 34 |
| *evm.model.Pm02.198* | Phosphatidylinositol 4-kinase gamma 3 |
| *evm.model.Pm02.199* | Dipeptidyl peptidase 8 |
| *evm.model.Pm02.200.2* | Calcineurin B-like protein 4 |
| *evm.model.Pm02.219* | Probable protein phosphatase 2C 76 |
| *evm.model.Pm02.220* | NA |
| *evm.model.Pm02.221* | Chaperone protein DnaJ |
| *evm.model.Pm02.222* | Probable trehalose-phosphate phosphatase 4 |
| *evm.model.Pm02.223* | Abscisic acid receptor PYL8 |
| *evm.model.Pm02.224* | NA |
| *evm.model.Pm02.225* | Polypyrimidine tract-binding protein homolog 2 |
| *evm.model.Pm02.226* | NA |
| *evm.model.Pm02.227* | Pentatricopeptide repeat-containing protein At2g13600 |
| *evm.model.Pm02.228* | Bidirectional sugar transporter SWEET3 |
| *evm.model.Pm02.385* | RHOMBOID-like protein 2 |
| *evm.model.Pm02.386* | Putative non-specific lipid-transfer protein 14 |
| *evm.model.Pm02.387* | Protein IQ-DOMAIN 14 |
| *evm.model.Pm02.388* | Protein FAR1-RELATED SEQUENCE 5 |
| *evm.model.Pm02.389* | Protein MEN-8 |
| *evm.model.Pm02.390* | Probable methyltransferase PMT16 |
| *evm.model.Pm02.391* | E3 ubiquitin ligase BIG BROTHER-related |
| *evm.model.Pm02.392* | F-box protein PP2-A13 |
| *evm.model.Pm02.393* | Probable transcriptional regulator SLK3 |
| *evm.model.Pm02.394* | Dihydrodipicolinate reductase-like protein CRR1, chloroplastic |
| *evm.model.Pm02.395* | BAG family molecular chaperone regulator 1 |
| *evm.model.Pm02.396* | MATE efflux family protein LAL5 |
| *evm.model.Pm02.397* | Serine/arginine-rich splicing factor RS40 |
| *evm.model.Pm02.398* | TraB domain-containing protein |
| *evm.model.Pm02.400* | Ethylene-responsive transcription factor ERF025 |
| *evm.model.Pm02.401* | Dehydration-responsive element-binding protein 1D |
| *evm.model.Pm02.7* | Glioma tumor suppressor candidate region gene 1 protein |
| *evm.model.Pm02.8* | NA |
| *evm.model.Pm02.9* | Origin of replication complex subunit 3 |
| *evm.model.Pm02.99* | Uncharacterized calcium-binding protein At1g02270 |
| *evm.model.Pm08.1751* | Costars family protein |
| *evm.model.Pm08.1752.1* | KH domain-containing protein At4g18375 |
| *evm.model.Pm08.1753* | Threonine--tRNA ligase, mitochondrial 1 |
| *evm.model.Pm08.1754* | Fructose-1,6-bisphosphatase, chloroplastic |
| *evm.model.Pm08.1755* | Protein EXECUTER 1, chloroplastic |
| *evm.model.Pm08.1756* | Beta-ureidopropionase |
| *evm.model.Pm08.1757* | NA |
| *evm.model.Pm08.1758* | General transcription factor 3C polypeptide 3 |
| *evm.model.Pm08.1759* | Uncharacterized protein At4g38062 |
| *evm.model.Pm08.1760* | Magnesium-dependent phosphatase 1 |
| *evm.model.Pm08.1761* | Periaxin |
| *evm.model.Pm08.1762* | NA |
| *evm.model.Pm08.1763* | Probable ubiquitin-like-specific protease 2B |

Note: ‘NA’ indicated Not Applicable.

**Supplementary Table 24 Selected genes in the KEGG pathways associated with plateau adaptability.**

| KEGG pathway | Gene ID | Function annotated by Swiss-Prot database |
| --- | --- | --- |
| DNA repair | *evm.model.Pm07.607* | Fanconi anemia group E protein |
|  | *evm.model.Pm01.2340* | Protein BCCIP homolog |
|  | *evm.model.Pm01.2341* | Protein BCCIP homolog |
|  | *evm.model.Pm01.2365* | ATP-dependent DNA helicase RecG |
| Inhibition cell death | *evm.model.Pm07.598* | Graves disease carrier protein |
|  | *evm.model.Pm05.1734* | Delta-1-pyrroline-5-carboxylate dehydrogenase 12A1 |
|  | *evm.model.Pm06.1723* | Protein CPR-5 |
|  | *evm.model.Pm01.3748* | Protein ROOT PRIMORDIUM DEFECTIVE 1 |
| Cold resistance | *evm.model.Pm07.600* | Serine acetyltransferase 5 |
|  | *evm.model.Pm07.603* | Probable acyl-[acyl-carrier-protein]--UDP-N-acetylglucosamine O-acyltransferase |
|  | *evm.model.Pm07.676* | Phospholipid:diacylglycerol acyltransferase 1 |
|  | *evm.model.Pm07.677* | Phospholipid:diacylglycerol acyltransferase 1 |
|  | *evm.model.Pm05.1686* | Germin-like protein subfamily T member 2 |
|  | *evm.model.Pm05.1688* | Germin-like protein subfamily T member 2 |
|  | *evm.model.Pm05.1689* | Germin-like protein subfamily T member 2 |
|  | *evm.model.Pm05.1691* | Germin-like protein subfamily T member 2 |
|  | *evm.model.Pm05.1735* | Probable WRKY transcription factor 72 |
|  | *evm.model.Pm02.386* | Putative non-specific lipid-transfer protein 14 |
|  | *evm.model.Pm02.401* | Dehydration-responsive element-binding protein 1D |
| UV light response | *evm.model.Pm01.3432* | Transcription factor HY5-like |
|  | *evm.model.Pm08.1755* | Protein EXECUTER 1, chloroplastic |
|  | *evm.model.Pm02.394* | Dihydrodipicolinate reductase-like protein CRR1, chloroplastic |
|  | *evm.model.Pm02.388* | Protein FAR1-RELATED SEQUENCE 5 |
